# When Does Treatment Memory Promote Glioma Growth? Feedback Placement and Schedule-Dependent Dormancy

**DOI:** 10.64898/2026.11.23.768612

**Authors:** Gabriel Feumi-Jantou, Naghmeh Akhavan

## Abstract

We study how treatment-history feedback changes phenotype composition and cumulative growth in a two-phenotype glioma model. Memory enters switching (M1), proliferative fitness (M2), or both (M3). An exact phase-resolved identity accounts for the observed schedule-dependent growth reversals, while an adjoint formula describes only the weak-feedback response. Across 2,000 sampled scenarios, 45 exact M1/base reversals occur, of which the first-order formula detects 13. A fitness-allocation counterexample shows that identical phenotype trajectories need not imply identical population growth. Multistart continuous fitting of noisy synthetic observations illustrates how uncertain baseline parameters complicate mechanism discrimination. A density-dependent extension shows that time-integrated growth retains information largely absent from saturated endpoint populations. These results concern phenomenological models with uncalibrated parameters, not clinical treatment predictions.

## 1 Introduction

Glioblastoma (GBM) is a highly aggressive primary brain tumor characterized by extensive cellular heterogeneity and poor clinical outcomes despite multimodal treatment [1, 2]. Even after therapy, slowly cycling or dormant cell subpopulations may survive and later re-enter proliferation [3–5], motivating mathematical descriptions of transitions between proliferating (*P*) and dormant (*D*) phenotypes [6–8]. Cellular dormancy refers here to reversible suppression of proliferation at the cell level, as distinct from tumor-mass dormancy in which proliferation and loss can balance at the population level.

Studies of temozolomide (TMZ)-treated glioblastoma have described treatment-associated dormancy, exit from dormancy after withdrawal, and transient slow-growing states during the acquisition of resistance [3, 5, 6, 9, 10]. Broader evidence from other cancer systems implicates reversible drug-tolerant states and treatment-associated reprogramming [11, 12]. These observations motivate, but do not directly validate, a mathematical memory law for glioma phenotype switching or growth.

Böttcher et al. [6] developed an evolutionary-game-theoretic model for two interacting phenotypes, in which current treatment reduces proliferative fitness while reversible phenotype conversion occurs at a constant rate. Their framework permits analysis of equilibria and alternating high/low treatment schedules, but it does not retain information about exposure once the current phenotype composition and treatment input have been specified. Thus populations with the same current state and treatment follow the same subsequent phenotype dynamics regardless of prior exposure.

The gap considered here is more specific than the absence of an additional state variable: it is not known, within this minimal evolutionary framework, whether persistent exposure effects acting on phenotype conversion can be distinguished from those acting directly on proliferative fitness. Even locally similar phenotype responses need not imply similar cumulative population growth. This leads to two questions: what dynamical signatures are generated by different memory placements, and which conclusions can be established from the equations rather than inferred from numerical trajectories alone?

We represent recent treatment history by a bounded state variable satisfying a first-order relaxation law. This simple exponential filter is a phenomenological approximation rather than a new memory operator or a claim about specific intracellular pathways. We compare three feedback hypotheses: memory-dependent asymmetric switching (M1); an additional, persistent proliferative-fitness cost (M2); and combined switching and fitness feedback (M3). Their selected signs and functional forms are exploratory modeling assumptions requiring biological validation.

Our main contributions are (i) the exact high/low phase decomposition of switching-memory growth, including numerical sign reversals; (ii) a finite-horizon adjoint formula for weak feedback together with an explicit demonstration of its finite-feedback limitations; (iii) a fitness-allocation counterexample establishing the ambiguity of phenotype-only data; and (iv) numerical comparisons of trajectory matching, treatment schedules, density-dependent population growth, and model discrimination with uncertain baseline parameters. The exponential filter itself is standard; our focus is how feedback placement and treatment timing alter the consequences of memory.

Related work has explicitly modeled therapy-associated transitions into and out of dormancy, including stochastic, treatment-dependent switching models with multiple drug protocols [13]. Accordingly, treatment-modulated phenotype switching is not itself unique to this study. Our narrower contribution is the placement-specific comparison within the Böttcher evolutionary-fitness formulation and its exact growth and output-signature identities. A recent computational review likewise identifies the difficulty of connecting temporally sampled dormancy observations to the underlying switching mechanisms [14]. Related work on treatment-induced versus spontaneous resistance and phenotypic plasticity has addressed similar mechanistic distinctions using mathematical and experimental approaches [15–18]. The first-order memory filter is closely related in mathematical structure to classical pharmacodynamic effect-compartment dynamics [19]; our normalized exposure state, however, is not a measured drug concentration or a validated effect compartment. The added contribution here is a controlled comparison of specific feedback allocations and their output implications, not the discovery of a new filtering or identifiability principle. The algebraic residual separation also assumes that treatment memory does not change dormant-cell fitness; with feedback in both phenotype fitnesses, two output residuals cannot identify all feedback channels without additional restrictions.

The study is theoretical and exploratory. Treatment cost and model time are nondimensional, the growth observable relies on identifying mean evolutionary fitness with an effective net growth rate, and the proposed feedback parameters are not fitted to longitudinal tumor data. Consequently, comparisons are interpreted as consequences of explicitly stated mathematical mechanisms rather than recommendations for clinical chemotherapy schedules.

## 2 Mathematical Model

We first recall the two-phenotype base model, then define treatment memory and its alternative placements in the evolutionary equations.

### 2.1 The base model

Böttcher et al. [6] considered two phenotype fractions *x*_*D*_ + *x*_*P*_ = 1 with payoff matrix (ordered *D, P*)

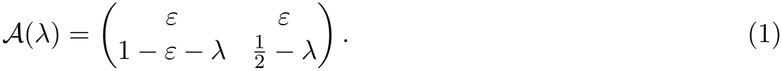

Here *ε* denotes dormant-cell fitness and *λ*(*t*) is a treatment-dependent fitness cost acting on proliferating cells. The phenotype fitness functions and population mean are

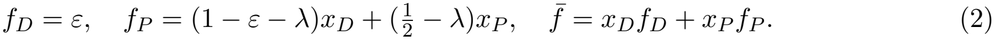

The original frequency dynamics are

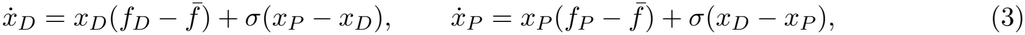

where the spontaneous conversion rates are symmetric and equal to *σ*. With *x* = *x*_*D*_ and *x*_*P*_ = 1 − *x*, define

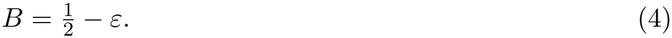

The scalar equation and mean fitness become

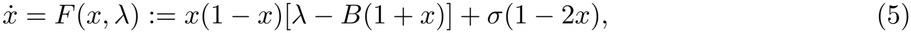

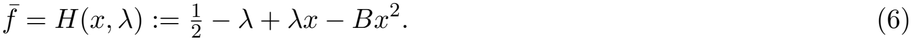

Because the phenotype fractions do not determine total population size, we additionally use the auxiliary growth relation

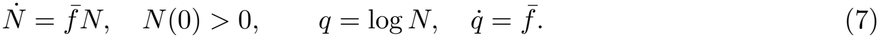

Equation (7) assumes mean evolutionary fitness represents an effective per-capita net growth rate; it is not an additional equation in the published phenotype-frequency system. Without resource constraints or a mapping to physical time, *N* is a normalized cumulative-growth observable, not a calibrated cell count or spatial tumor volume. The growth rate *G*(*T*) = (*q*(*T*) − *q*(0))*/T* and relative population ratios are therefore preferable to absolute numerical values of *N*.

If the payoffs *f*_*D*_ and *f*_*P*_ are interpreted as absolute net per-capita growth rates, the frequency and total-growth equations are equivalent to the compartment system

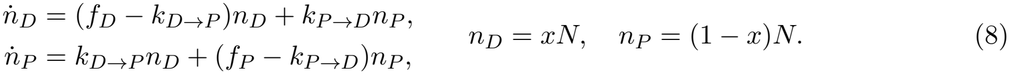

This derivation makes the additional assumption explicit: *f*_*D*_ = *ε* = 0 represents zero *net* birth-minus-death rate of dormant cells before switching, not biological absence of both division and death. By contrast, replicator fractions alone identify fitness differences but are invariant to adding the same function of time to both fitnesses; absolute population growth is not invariant to that shift.

### 2.2 Exponentially Filtered Treatment History

Let *λ*_*H*_ *>* 0 and 0 ≤ *λ*_*L*_ *< λ*_*H*_ denote high and low treatment fitness costs. The protocol *λ*(*t*) may alternate between these levels or contain treatment-free intervals. We normalize *u*(*t*) = *λ*(*t*)*/λ*_*H*_ ∈ [0, 1] and introduce

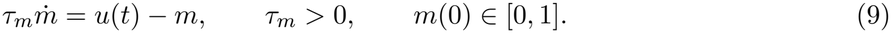

For prescribed exposure,

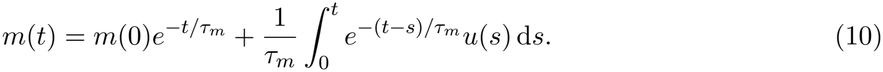

Thus more recent exposures receive greater weight, and the history state relaxes following treatment withdrawal. This is a minimal phenomenological law rather than a validated pharmacokinetic or molecular model.

### 2.3 M1: Memory-Dependent Phenotype Switching

Inspired by evidence that entry into and exit from TMZ-associated dormancy can be regulated [3], Model M1 assumes

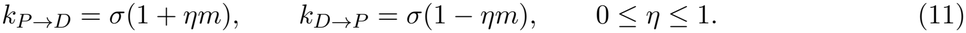

The rates are nonnegative and sum to 2*σ*. This exactly balanced alteration is a parsimonious structural choice, not an experimentally established equality between the response of dormancy entry and exit. M1 retains the original fitness functions:

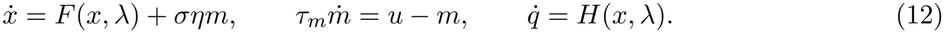

At *η* = 0 the (*x, q*) evolution coincides with the base model.

### 2.4 M2 and M3: Fitness and Combined Memory Feedback

Motivated by persistent, non-genetic changes in treatment responses [11, 12], Model M2 assumes an additional fitness penalty on the proliferating population:

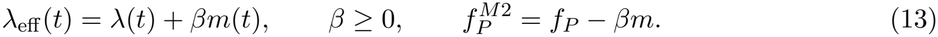

The symmetric phenotype conversion rates remain unchanged. Model M3 combines the M1 switching asymmetry with the M2 penalty. With indicators (*I*_*S*_, *I*_*F*_) = (1, 0), (0, 1), (1, 1) for M1, M2, M3 respectively, the unified system reads

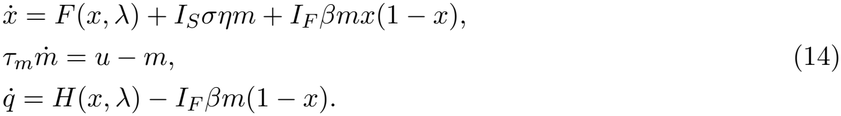

This schematic emphasizes that changing phenotype frequencies and changing absolute net population growth are distinct modeling operations. A single shared memory state represents the exposure history of the aggregate population; individual cells do not carry separate memory variables.

The assumption of persistent fitness suppression is motivated by experimental evidence that glioblastoma cells surviving temozolomide exposure can exhibit altered proliferative behavior during post-treatment recovery. In particular, Silva et al. [20] reported differences in the regrowth kinetics of treated and untreated glioblastoma cell populations. These observations provide a biological motivation for incorporating treatment-history-dependent fitness effects, although the specific residual fitness term *βm* has not been directly calibrated from these experiments. Crucially, M2 holds dormant-cell fitness fixed: Δ_*D*_ = 0 and Δ_*P*_ = −*βm*. This restriction is part of the model definition and will be tested below against an alternative allocation of the same *fitness difference*.

## 3 Mathematical Properties

We first establish admissible states and constant-treatment stability, then describe the local response to oscillatory exposure. These elementary properties support the mechanism comparisons developed in Section 4.

### Proposition 3.1

(**Forward invariance and well-posedness**). *If σ* ≥ 0, *τ*_*m*_ *>* 0, 0 ≤ *η* ≤ 1, *β* ≥ 0, *u* ∈ [0, 1], *x*(0), *m*(0) ∈ [0, 1], *and N* (0) *>* 0, *then for bounded piecewise-continuous treatment the unique solution exists on each finite time interval and satisfies x, m* ∈ [0, 1] *and N >* 0.

*Proof*. At *x* = 0 the vector field in Eq. (14) is *σ*(1 + *I*_*S*_*ηm*) ≥ 0; at *x* = 1 it is −*σ*(1 − *I*_*S*_*ηm*) ≤ 0. The memory derivative is nonnegative at *m* = 0 and nonpositive at *m* = 1. These inward-pointing inequalities give forward invariance. The vector field is locally Lipschitz in the states on constant-input intervals; boundedness of *x, m* and of 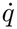 gives continuation over finite intervals. Since 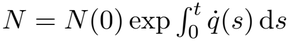, positivity follows. Solutions remain continuous across treatment switches.

### 3.1 Constant Exposure and Local Stability

For *λ*(*t*) = *λ*_0_, the memory equilibrium is *m*_*\**_ = *λ*_0_*/λ*_*H*_. The phenotype equilibria satisfy

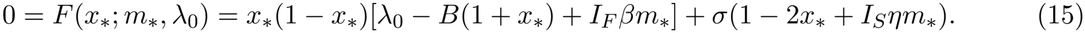

For *σ >* 0, the right-hand side is positive at *x* = 0 and nonpositive at *x* = 1, so a root exists in [0, 1]. Its number need not be unique without further hypotheses. The Jacobian of the (*x, m*) dynamics at equilibrium is

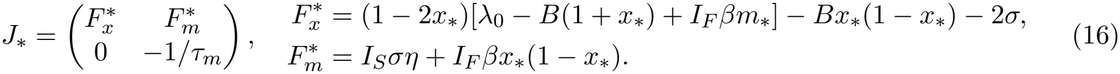

A hyperbolic phenotype-memory equilibrium is locally asymptotically stable precisely when 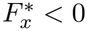. The equilibrium location is independent of *τ*_*m*_, whereas transient relaxation is not. The auxiliary growth variable need not be stationary at phenotype-memory equilibrium.

A numerical examination of the constant-treatment equilibrium equations identified a single biologically admissible equilibrium for each model at all 1,001 sampled treatment levels under the nominal parameters. Although this does not establish uniqueness analytically, the numerical results provide additional information about the equilibrium structure in the investigated parameter regime (Supplementary Section A.3).

### 3.2 Small Periodic Perturbations

For 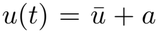 sin(*ωt*) with 0 ≤ *u* ≤ 1, the memory equation has the unique attracting periodic solution

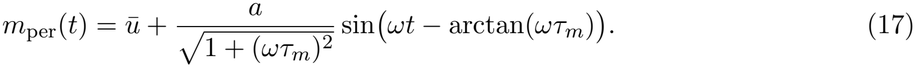

Hence rapidly varying treatment is attenuated in the history state. The nonlinear phenotype variable also responds directly to current exposure; its frequency response cannot be deduced from Eq. (17) alone.

#### Proposition 3.2

(**Local periodic response**). *Suppose* 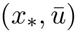 *is an equilibrium under constant* 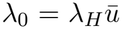 *with*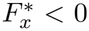. *For each fixed ω >* 0 *and sufficiently small a, the periodically forced* (*x, m*) *system has a locally unique stable periodic orbit near*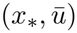. *Its first-order component is*

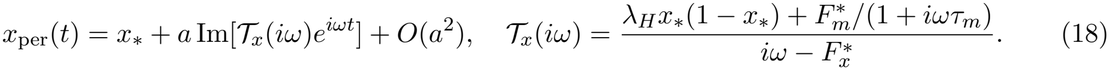

*Proof*. At *a* = 0, both eigenvalues 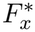 and −1*/τ*_*m*_ are strictly negative. The implicit-function theorem on the period map and continuity of Floquet multipliers provide persistence and stability of the nearby periodic orbit. The first variation obeys 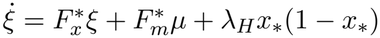 and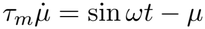, yielding Eq. (18) by harmonic substitution.

The result characterizes the local response to small-amplitude periodic forcing. It does not, by itself, establish an optimal treatment period for finite-amplitude, alternating high/low treatment protocols.

## 4 Analytical Results: Comparison of Memory Mechanisms and Periodic Treatment

The numerical figures below motivate three analytical questions: whether dormant-cell fractions can be ordered; how the different feedback pathways contribute to cumulative population growth; and which features of periodic exposure survive fast-switching limits. We additionally ask whether the two memory pathways have distinct mathematical signatures in joint phenotype and growth observations. The base model, defined by Eqs. (5) and (7), and the memory-dependent Models M1 and M2, given by Eq. (12) and Eq. (14) with (*I*_*S*_, *I*_*F*_) = (0, 1), respectively, are compared under the same treatment protocol *λ*(*t*), with identical initial dormant-cell fractions *x*(0) = *x*_0_ and total modeled populations *N* (0) = *N*_0_ *>* 0. The memory-dependent models also share the same initial memory state *m*(0) = *m*_0_, governed by Eq. (9). Throughout this section, 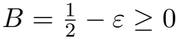, *σ >* 0, 0 ≤ *η* ≤ 1, *β* ≥ 0, and *τ*_*m*_ *>* 0. For clarity let (*x*_base_, *q*_base_), (*x*_1_, *q*_1_), and (*x*_2_, *q*_2_) refer to the base, M1 and M2 states, respectively:

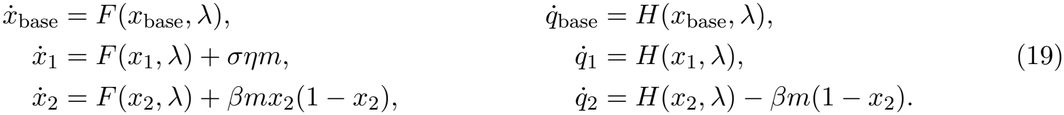

The invariant-state result in Proposition 3.1 applies to each system.

### 4.1 Ordering of dormant-cell fractions

We first examine how the placement of treatment memory affects the evolution of the dormant-cell fraction. Although both M1 and M2 introduce memory-dependent contributions that favor dormancy, they act through different mechanisms: M1 modifies phenotype-switching rates, whereas M2 alters the relative fitness of proliferating cells. We therefore investigate whether these mechanisms imply a consistent ordering of dormant-cell fractions under identical treatment histories and initial conditions. The following proposition establishes such an ordering under explicit conditions on the memory-feedback parameters.

#### Proposition 4.1

(**Phenotype comparison**). *For equal initial dormant fractions and common exposure, x*_1_(*t*) ≥ *x*_base_(*t*) *at all finite times. If in addition* 0 ≤ *β* ≤ 4*ση, then x*_1_(*t*) ≥ *x*_2_(*t*) *at all finite times*.

*Proof*. The difference *z* = *x*_1_−*x*_base_ satisfies *ż* = *a*(*t*)*z* + *σηm*(*t*), where *a*(*t*) is a bounded divided difference of *F* in its first argument. Variation of constants gives 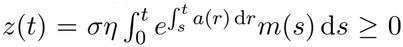. For M1 versus M2, at any common *x* ∈ [0, 1] their vector-field difference is *m*[*ση* − *βx*(1 − *x*)] ≥ *m*(*ση* − *β/*4) ≥ 0. Scalar comparison completes the proof.

The nominal midpoint calibration, *β* = 4*ση*, satisfies the sufficient condition in Proposition 4.1. However, for a general reference fraction, the local matching condition *β* = *ση/*[*x*_ref_ (1 − *x*_ref_)] yields *β* ≥ 4*ση*, with equality only when *x*_ref_ = 1*/*2 (for *ση >* 0). Consequently, the proposition does not establish the M1–M2 dormant-fraction ordering for the non-midpoint calibrations used in the parameter sweeps. The corresponding numerical observations must therefore be distinguished from the analytical comparison result.

### 4.2 Exact decomposition of relative population growth

The phenotype comparison alone does not order the total modeled populations because *H*(*x, λ*) need not be increasing in *x*. With *z* = *x*_1_ − *x*_2_, the polynomial identity

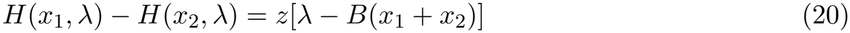

yields an exact decomposition.

#### Proposition 4.2

(**Mechanism-Resolved Cumulative Growth**). *For common initial populations and every finite horizon T*,

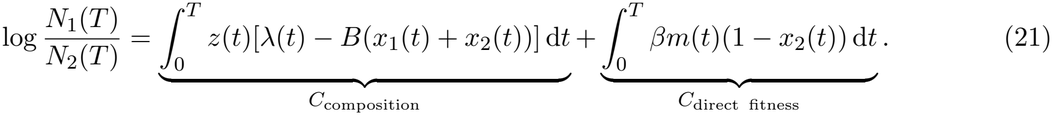

*Proof*. Subtract the two *q* equations in Eq. (19), apply Eq. (20), and integrate.

The second integrand is nonnegative by construction, whereas the first may have either sign. Thus an opposite-sign response between M1 and M2 is not an assumption-free property of memory; a direct negative fitness feedback is explicitly present in M2.

For the nominal three-cycle treatment protocol, we numerically evaluated the two contributions in Eq. (21) by integrating over each treatment phase. The resulting decomposition, reported in Table 2, gives *C*_composition_ = 0.240676 and *C*_direct fitness_ = 1.030704, with a total logarithmic population difference of log[*N*_1_(*T*)*/N*_2_(*T*)] = 1.271380. Thus, approximately 18.9% of this difference is associated with phenotype composition, while 81.1% arises from the direct fitness-memory term in M2.

**Table 1:**
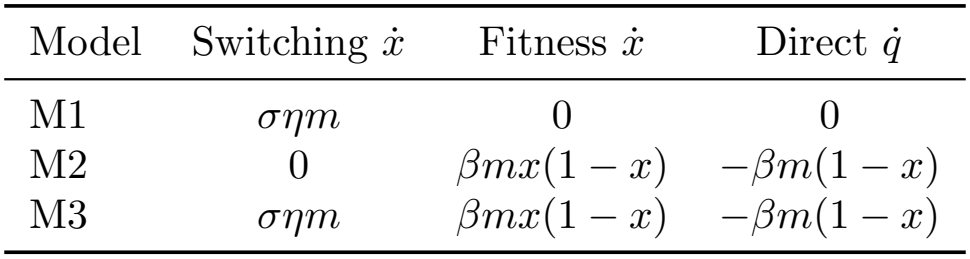
Memory-feedback contributions in M1–M3. Additive changes to the dormant-fraction equation and to the modeled logarithmic population-growth equation.

| Model | Switching ẋ | Fitness ẋ | Direct q̇ |
|---|---|---|---|
| M1 | σηm | 0 | 0 |
| M2 | 0 | βmx(1 − x) | −βm(1 − x) |
| M3 | σηm | βmx(1 − x) | −βm(1 − x) |

**Table 2:** Numerical decomposition of nominal M1–M2 log-population separation. Summed contributions over all intervals of each treatment level in the three-cycle schedule. Values are obtained by phase-segmented numerical integration of Eq. (21) for the protocol described in Supplementary Table 6.

| Exposure phase | Composition | Direct M2 penalty | Sum |
|---|---|---|---|
| Untreated | 0 | 0 | 0 |
| High treatment | +0.306133 | +0.378044 | +0.684177 |
| Low treatment | −0.065457 | +0.652661 | +0.587203 |
| All phases | +0.240676 | +1.030704 | +1.271380 |

These two terms are an exact algebraic accounting of the specified M1–M2 trajectories, not independent causal effects. In particular, the composition term also depends on the M2 fitness parameter through the induced trajectory *x*_2_(*t*). The 18.9%/81.1% shares cannot be transferred unchanged to a model that allocates treatment-related fitness differently between phenotypes.

The biological relevance of this decomposition is supported by experimental and computational studies of glioblastoma that distinguish changes in cellular states from changes in population growth dynamics. In particular, Larsson et al. [21] used mathematical modeling of lineage-tracing data to quantify cell-state transitions and state-specific growth rates, while Adamski et al. [3] demonstrated treatment-associated dormancy and subsequent reactivation. Silva et al. [20] further reported altered regrowth kinetics in glioblastoma cells following temozolomide exposure. Motivated by these observations, our exact decomposition quantifies the contributions of phenotype composition and direct fitness feedback to the difference in cumulative logarithmic population growth between M1 and M2, providing a framework for investigating how distinct treatment-memory mechanisms influence population dynamics.

### 4.3 Schedule-dependent sign reversal and the adjoint value of dormancy

The phenotype comparison in Proposition 4.1 implies *z*_*b*_ = *x*_1_ − *x*_base_ ≥ 0, but it does not fix the sign of cumulative growth. Subtracting the base and M1 logarithmic-growth equations gives an exact identity.

#### Proposition 4.3

(**Exact phase-resolved growth comparison**)

*For identical initial populations under any common treatment history*,

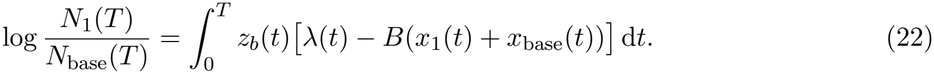

*For an initial untreated phase with m*(0) = 0 *followed by high- and low-treatment intervals, write the corresponding integrals as C*_*H*_ *and C*_*L*_. *The untreated contribution vanishes, so N*_1_(*T*) *< N*_base_(*T*) *if and only if C*_*H*_ + *C*_*L*_ *<* 0. *In particular, if λ*_*H*_ ≥ 2*B, then C*_*H*_ ≥ 0 *and* −*C*_*L*_ *> C*_*H*_ *is necessary and sufficient for reversal*.

*Proof*. The identity follows from 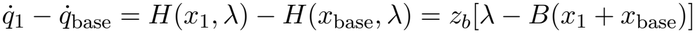 and equal initial populations. Before the first exposure the memory is zero and both phenotype systems have identical initial values; uniqueness implies *z*_*b*_ = 0. On high-treatment phases *λ*_*H*_ − *B*(*x*_1_ + *x*_base_) ≥ *λ*_*H*_ − 2*B* ≥ 0.

The sign is determined by the interaction of phenotype differences and current treatment, rather than by dormancy alone. Strong exposure can generate a positive composition contribution, whereas persistent differences during near-withdrawal intervals can make the low-treatment contribution negative and larger in magnitude. This exact criterion does not require small memory feedback, but it is trajectory-dependent rather than a parameter-only schedule-optimization theorem.

We next derive a general first-order formula that quantifies the cumulative-growth effect of a weak memory-dependent perturbation. The perturbation parameter *ρ* below is independent of the dormant-fitness parameter *ε*.

#### Proposition 4.4

(**Adjoint linear response of cumulative growth**). *Let* (*x*_*b*_, *m, q*_*b*_) *be the base trajectory under a bounded piecewise-continuous treatment protocol on* [0, *T*], *with F, H continuously differentiable in x. Consider the perturbed equations*

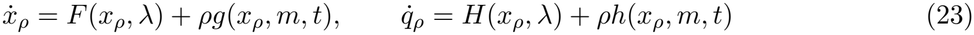

*with identical initial states and sufficiently smooth bounded perturbations g, h. Define ψ backward by*

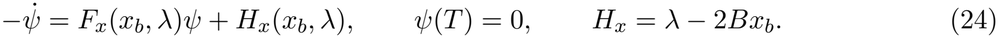

*Then for fixed T as ρ* → 0,

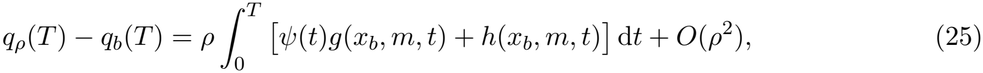

*where*

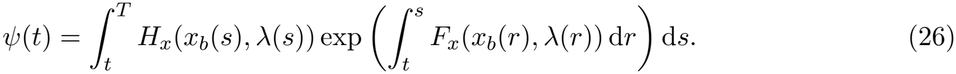

*Proof*. Set *ξ* = *∂*_*ρ*_*x*_*ρ*_|_*ρ*=0_ and *ζ* = *∂*_*ρ*_*q*_*ρ*_|_*ρ*=0_. Differentiating the equations gives 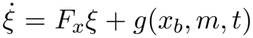, *ξ*(0) = 0, and 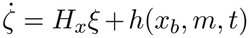, *ζ*(0) = 0. By Eq. (24),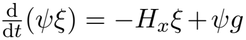. Integrating and using *ψ*(*T*) = 0 and *ξ*(0) = 0 yields 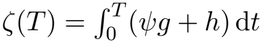. Smooth dependence on *ρ* over the finite horizon gives the remainder. Variation of constants in the backward adjoint equation gives Eq. (26).

The adjoint *ψ*(*t*) measures the first-order change in final log population caused by an infinitesimal increase in dormant fraction at time *t* propagated through the base dynamics. It may be positive or negative, because the future fitness benefit of dormancy depends on subsequent treatment and reactivation. For the three feedback placements and a memory benefit +*βm* acting on dormant fitness, Eq. (25) yields

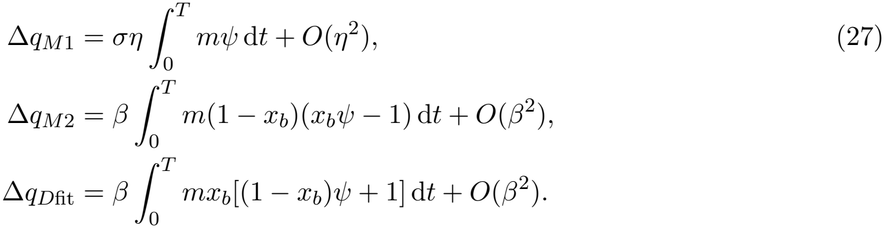

The response of M3 is the sum of its switching and fitness first variations, up to quadratic terms. These formulas hold for weak feedback on a fixed horizon; they do not provide exact signs or small errors for arbitrary *η* or *β*. In particular, the *O*(*η*^2^) remainder need not be negligible at the nominal *η* = 0.5: the sign of the first variation is not a reliable finite-feedback reversal classifier. We quantify this discrepancy against all 2,000 sampled protocols in Section 5.8.

### 4.4 Exact Dynamical Signatures of Memory Placement

An important question arising from the comparison of memory mechanisms is whether similar phenotype trajectories can conceal distinct feedback pathways. Although M1 and M2 can produce comparable changes in the dormant-cell fraction, their memory-dependent contributions enter different components of the governing equations. We therefore examine whether these contributions can be distinguished using joint observations of phenotype composition and cumulative population growth.

#### Proposition 4.5

(**Separation of Switching and Fitness Feedback**)

*Suppose F and H in Eqs*. (5)*–*(6) *are known, and let a differentiable trajectory* (*x*(*t*), *q*(*t*)) *satisfy Eq*. (14). *Define the model residuals*

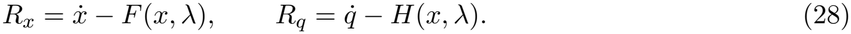

*Then the combinations*

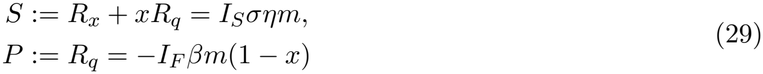

*separate the contributions of memory-dependent switching and direct fitness feedback exactly. When m >* 0, 0 ≤ *x <* 1, *and σ >* 0, *the active feedback amplitudes are recoverable as*

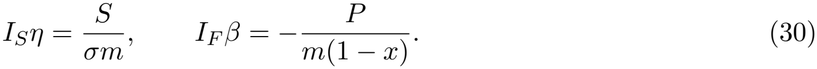

*Proof*. Substituting Eq. (14) into Eq. (28) gives

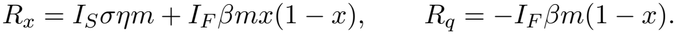

Consequently, the fitness-memory contribution to *R*_*x*_ cancels with *xR*_*q*_, yielding *S* = *I*_*S*_*σηm*, while *P* = *R*_*q*_ = −*I*_*F*_ *βm*(1 − *x*).

The resulting residuals provide distinct signatures of the memory mechanisms. For M1, memory contributes exclusively to the switching signature, *S* = *σηm*, while *P* = 0. For M2, *S* = 0 and *P* = −*βm*(1 − *x*), whereas M3 may exhibit nonzero contributions in both channels.

Supplementary Figure S3 illustrates these mechanism-resolved signatures for M1 and M2 under the nominal three-cycle treatment protocol and locally matched memory-feedback parameters. Panels (A) and (B) compare the dormant-cell fractions and logarithmic population trajectories, respectively. Although the two models exhibit similar phenotype responses, their cumulative population-growth predictions differ. Panels (C) and (D) show that the residuals distinguish the mechanisms: M1 produces a nonzero switching signature with a vanishing fitness signature, whereas M2 exhibits the opposite pattern. The plot illustrates the identities from Proposition 4.5 using model-evaluated vector fields; it is not an independent statistical test or experimental validation.

The separation established above is an exact structural property of the proposed mathematical framework under known base-model parameters, known treatment history, and differentiable observations. It does not, by itself, guarantee that the mechanisms can be reliably distinguished from experimental data. In particular, sparse or noisy measurements may complicate estimation of the required derivatives, while uncertainty in the baseline parameters or memory timescale may confound the inferred feedback contributions. Practical mechanism identification would therefore require appropriate estimation methods, such as integral formulations or direct fitting of the dynamical models to longitudinal observations.

The separation is *conditional* on a phenotype-specific fitness restriction. For arbitrary additional net-fitness terms Δ_*D*_(*t*) and Δ_*P*_ (*t*) and switching feedback *s*(*t*), the generalized equations give

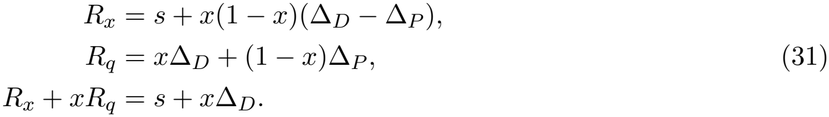

Thus a memory-dependent change in dormant-cell fitness contaminates the nominal switching signature *S*; with three unknown feedback functions (*s*, Δ_*D*_, Δ_*P*_) but only two observed residuals, the individual channels are not generally identifiable. Setting Δ_*D*_ = 0 restores Proposition 4.5. Moreover, with noise-less, continuously observed *x*(*t*) and known model parameters, M1 and M2 can already be distinguished generically from the differing *x*-dependences of their phenotype residuals. Joint observations improve information under noise; they are not mathematically necessary for ideal model-class discrimination.

### 4.5 Periodic Memory and Rapid Switching

For a repeated treatment cycle of length *T*_*c*_ *>* 0 with high-treatment fraction *d* ∈ (0, 1) and low normalized level *ℓ* = *λ*_*L*_*/λ*_*H*_, define 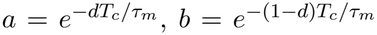. The memory return map is a contraction of slope *ab <* 1 and therefore has a unique globally attracting *T*_*c*_-periodic solution. Its minimum *m*_−_, maximum *m*_+_, mean, and amplitude are

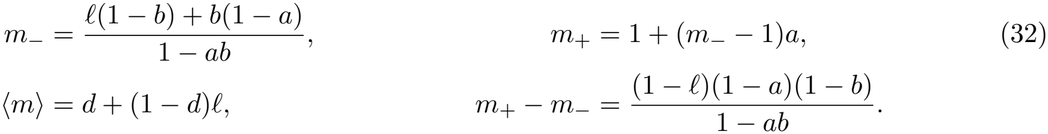

The mean is fixed by average exposure, whereas the amplitude varies with the cycle period relative to *τ*_*m*_. The phenotype dynamics also include instantaneous treatment terms, so memory amplitude alone does not order mean population growth across cycle periods.

#### Theorem 4.6

(**Finite-horizon fast-switching averaging**). *Fix a finite horizon T, τ*_*m*_ *>* 0, *bounded admissible parameters, and a high-treatment duty fraction d. Let T*_*c*_ = *T/n and apply n complete high/low cycles with common initial states. Let the averaged system have constant* 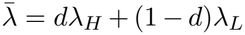 *and* 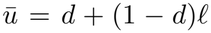. *For each model, there exists a constant C*_*T*_ *independent of n such that the maximum difference between the periodic-forcing solution* 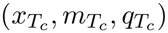 *and the averaged solution* 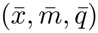 *is bounded by C*_*T*_ *T*_*c*_ *on* [0, *T*].

*Proof*. Write 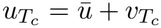 and 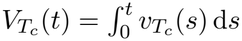. Each complete cycle has zero mean deviation, so 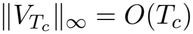. The state vector fields depend affinely on treatment input and have uniformly bounded state derivatives on the invariant state set over fixed *T*. Integration by parts in 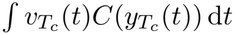, followed by subtraction of the integral equations and Gronwall’s inequality, gives an *O*(*T*_*c*_) bound uniformly on [0, *T*]. The logarithmic growth statement follows because *q* = log *N* is a state component.

Theorem 4.6 establishes convergence to the averaged dynamics as *T*_*c*_ → 0 on any fixed finite time horizon. The constant *C*_*T*_ may depend on *T*, so the estimate does not imply uniform convergence over arbitrarily long times. Moreover, the averaging result does not establish a monotonic relationship between treatment-cycle period and cumulative population growth, nor does it identify an optimal treatment period.

#### Corollary 4.7

(**Weak-feedback fast-switching equilibrium criterion**). *Consider the averaged model obtained in the vanishing-period limit, with constant input* 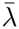 *and* 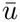, *and the corresponding base phenotype equilibrium* 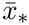 *is locally asymptotically stable, with* 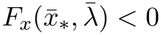. *For weak switching feedback η* ↓ 0, *the equilibrium mean log-growth shift satisfies*

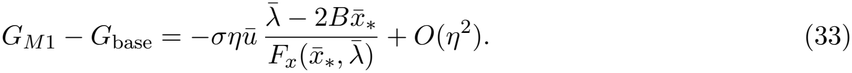

*Thus, when* 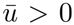 *and* 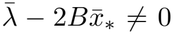, *the sign of the leading response equals that of*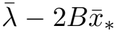. *If* 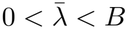 *stays away from B and σ is small, the approximation* 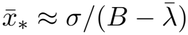 *gives the indicative inequality* 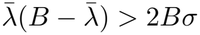 *for a positive leading response*.

*Proof*. At constant averaged input, 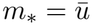 the equilibrium equation is 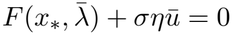. The implicit-function theorem gives 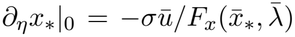. Differentiating 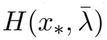 gives the stated first-order expression. The small-*σ* estimate follows by solving the equilibrium equation near 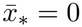 for 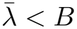.

The corollary concerns the averaged equilibrium and weak feedback, not every finite-period schedule. Neither it nor the finite-horizon averaging theorem establishes that all observed reversals require slow switching. A numerical averaged-equilibrium scan is reported in Supplementary Section A.2.

## 5 Numerical Results

We study how treatment-memory feedback changes phenotype composition and normalized cumulative growth, emphasizing which effects depend on the feedback mechanism. Unless stated otherwise, all simulations use the initial conditions and parameters in Supplementary Table 5; the treatment protocols are recorded in Supplementary Table 6. Population figures compare model-dependent growth indices and should not be read as clinical tumor-size forecasts. The original published baseline dynamics [6] are denoted the base model throughout this section.

### 5.1 Switching Memory changes phenotype composition and accumulated growth

M1 and the base model are compared under an identical three-cycle high/low protocol (Figure 1). In high-treatment intervals the dormant fraction increases, while the proliferative fraction recovers during low treatment. Since M1 retains the base fitness functions, its changes to mean population fitness arise indirectly through the altered phenotype composition (panels A and B). The exposure memory rises during high treatment and partially relaxes during low phases (panel D). The associated small differences in fitness accumulate over successive cycles (panels C and E). At *T* = 220, the nominal ratio *N*_1_*/N*_base_ ≃ 1.5971 corresponds to log(*N*_1_*/N*_base_) ≃ 0.4682, or 59.71% greater *total modeled population*.

**Figure 1:**
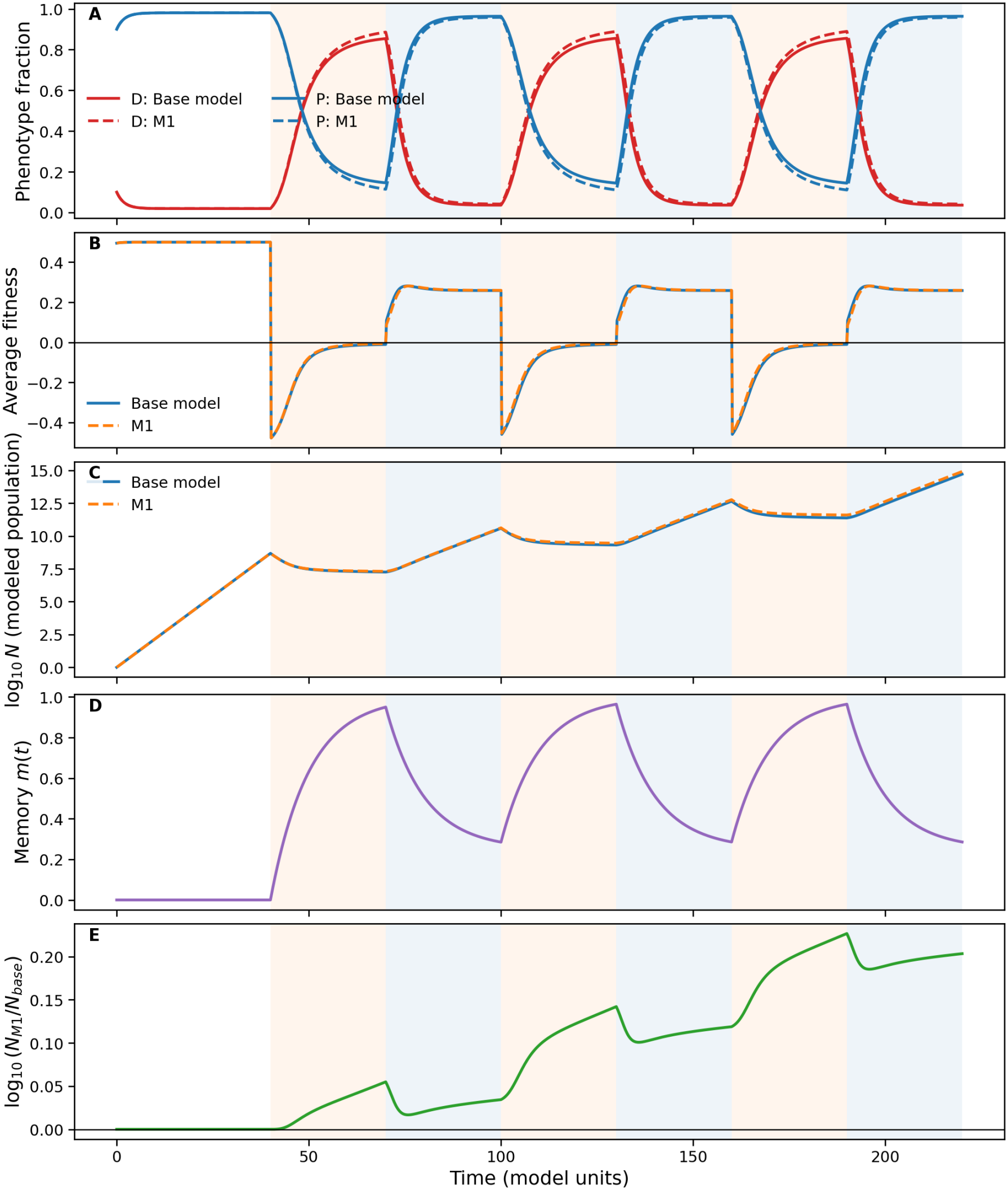
Memory-dependent switching alters composition and cumulative modeled growth. The base model and M1 receive 40 untreated time units followed by three high/low cycles with 30 units at *λ*_*H*_ = 1 and 30 at *λ*_*L*_ = 0.25 per cycle. Parameters: *ε* = 0, *σ* = 0.01, *η* = 0.5, *τ*_*m*_ = 10; *x*(0) = 0.1, *m*(0) = 0, *N* (0) = 1. (A) Dormant and proliferating fractions; (B) mean fitness; (C) log_10_ *N*; (D) exposure memory; (E) log_10_(*N*_1_*/N*_base_). The final modeled population is approximately 59.7% higher under M1.

This is not a claim about the number of dormant cells alone, nor a universal consequence of switching memory. The nominal *G* difference is 0.468195*/*220 ≈0.002128 per model-time unit, compared with the base rate *G*_base_ ≈0.153947, a relative rate increase of about 1.38%. Endpoint ratios accumulate over the long uncalibrated horizon and should not be interpreted as large instantaneous growth-rate changes.

### 5.2 Withdrawal exposes history dependence

A high-treatment pulse from *t* = 20 to *t* = 45 increases the dormant fraction in both models, followed by recovery after withdrawal (Figure 2A). The M1–base dormant-fraction difference temporarily persists after the pulse while *m* decays (panel C). To isolate retained history, two M1 trajectories start with the same *x*(0) = 0.8 and treatment *u* = 0, but with *m*(0) = 0 or 1 (panel B).

**Figure 2:**
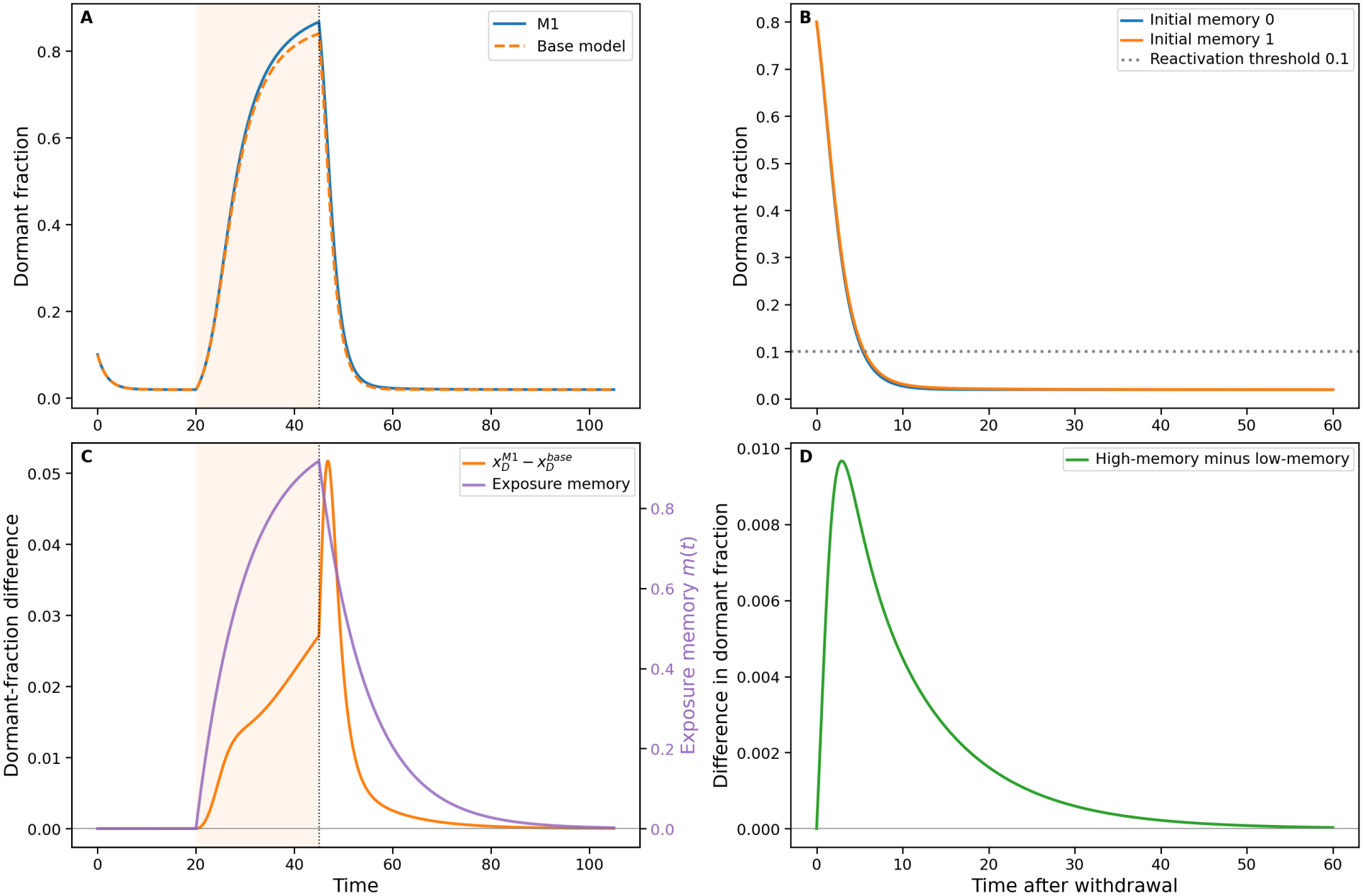
History-dependent dormant-cell reactivation after treatment withdrawal. (A) M1 and base dormant fractions under a pulse 20 ≤ *t* ≤ 45 and subsequent withdrawal; (B) equal initial dormant fractions *x*(0) = 0.8 under *u* = 0 but different memory states *m*(0) = 0, 1, with the dotted *x* = 0.1 threshold; (C) M1–base dormant-fraction difference and memory decay; (D) high-memory minus low-memory dormant fractions in the controlled comparison. Threshold crossing occurs at approximately 5.368 and 5.559 time units.

Defining reactivation operationally as the first crossing of *x* = 0.1, the times are approximately 5.368 and 5.559, a 0.191 model-time delay for the higher-memory state. The difference subsequently vanishes as memory relaxes (panel D). The *x* = 0.1 threshold is a numerical convention, not a biological criterion validated for GBM. The augmented (*x, m*) system remains Markovian even though observations of *x* alone omit the memory needed for prediction.

### 5.3 Sensitivity to memory strength and relaxation

We vary *η* ∈ [0, 1] and *τ*_*m*_ ∈ [1, 100] under the reference protocol, quantifying the final effect by

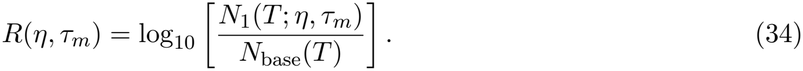

Figure 3 shows *R* = 0 at *η* = 0 (recovery of the base phenotype dynamics). Across the displayed parameter map, increasing *η* generally increases *R*, whereas the variation with *τ*_*m*_ is not uniformly monotonic. The indicated nominal point is (*η, τ*_*m*_) = (0.5, 10). Thus stronger memory feedback and longer memory persistence are not interchangeable; their influence depends on the imposed treatment waveform.

**Figure 3:**
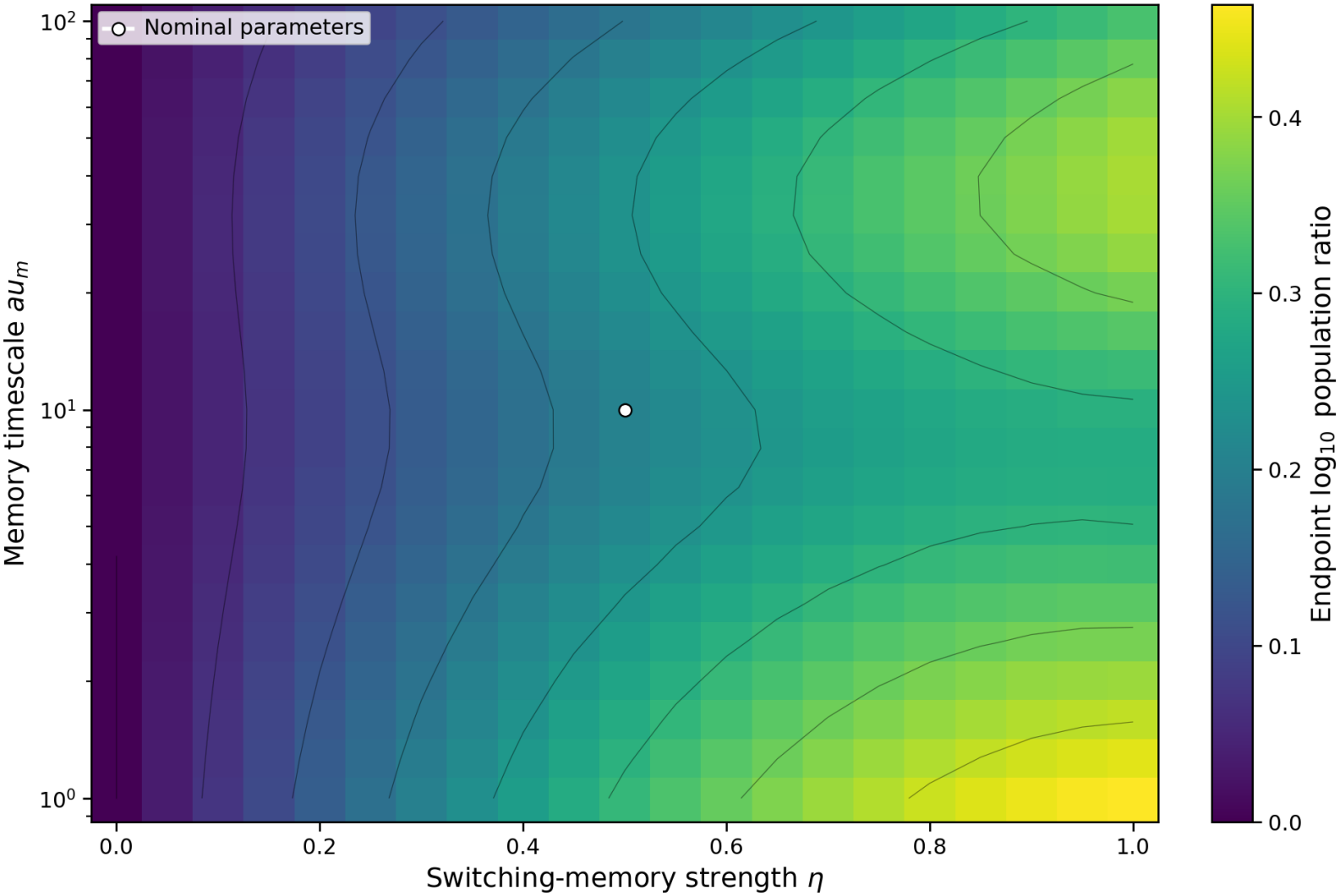
Sensitivity of final modeled population to switching-memory strength and timescale. Heat map of *R* = log_10_(*N*_1_*/N*_base_) for *η* ∈ [0, 1], *τ*_*m*_ ∈ [1, 100] (logarithmic vertical axis). Contours indicate equal *R*, and the white circle marks nominal (0.5, 10). Positive values indicate a larger final modeled population in M1 under this protocol.

### 5.4 Treatment-cycle period and memory persistence

The local sinusoidal response formula does not predict the nonlinear response under piecewise-constant high/low therapy. We therefore vary the complete treatment period *T*_*c*_ over a fixed horizon *T* = 240 with a 50% high/50% low duty cycle (*λ*_*H*_ = 1, *λ*_*L*_ = 0.25). All 14 simulated periods, *T*_*c*_ ∈ {4, 6, 8, 10, 12, 16, 20, 24, 30, 40, 48, 60, 80, 120}, divide 240 exactly. Each protocol therefore assigns exactly 120 time units to each treatment level, with cumulative exposure cost 150. The revised heatmap is clipped to the simulated period range 4 ≤ *T*_*c*_ ≤ 120. The comparison uses finite-horizon mean logarithmic growth

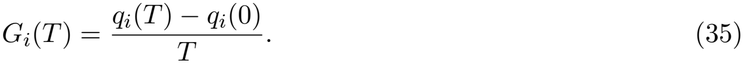

Figure 4A shows increasing *G* as cycle period grows over the sampled grid for both the base model and M1. M1 remains slightly above the base growth rate in the tested conditions. The heatmap (panel B) shows positive *G*_1_ − *G*_base_, with larger differences generally at shorter periods and a non-monotone dependence on memory relaxation. This is a change in *magnitude* of growth, not a reversal of the ranking of cycle periods.

**Figure 4:**
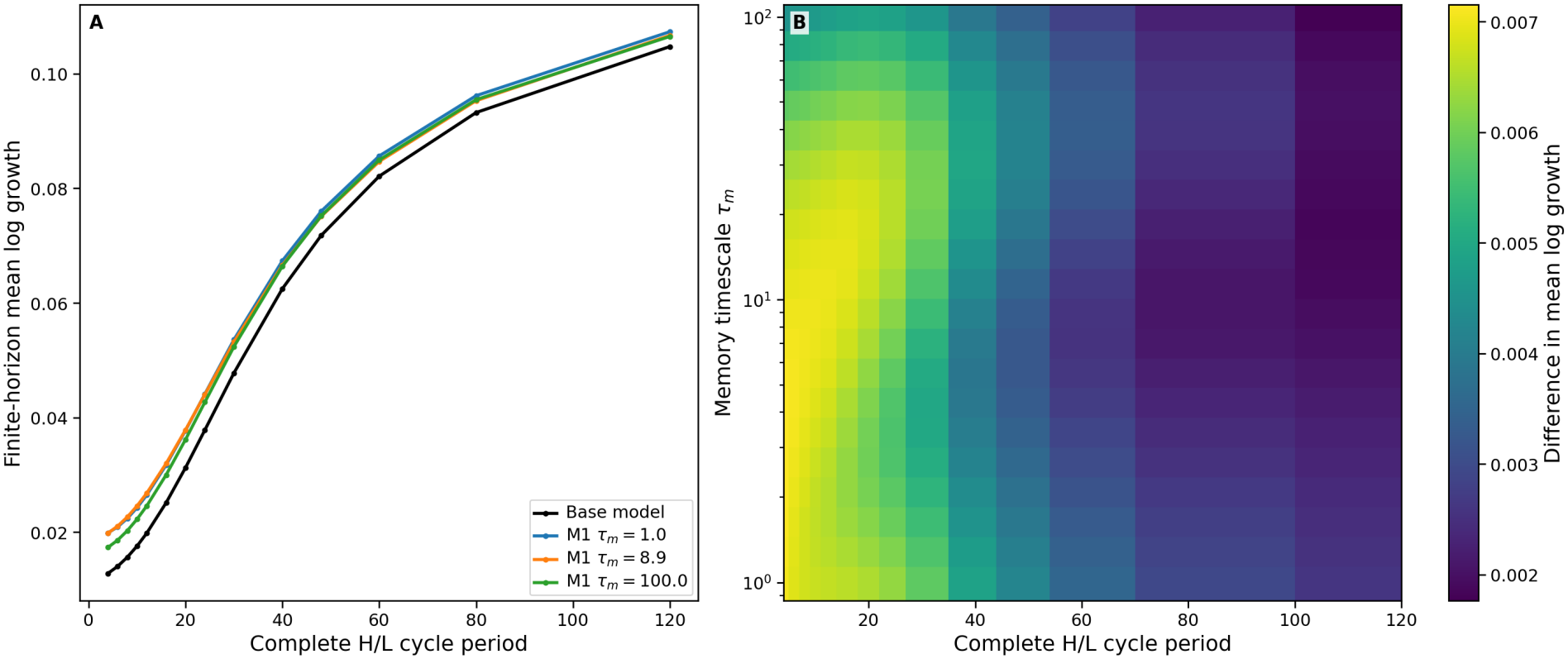
Treatment-period and memory-timescale effects on finite-horizon growth. (A) Mean logarithmic growth versus complete-cycle period, comparing the base model and M1 for representative *τ*_*m*_ ≈1, 8.9, 100. (B) Difference *G*_1_ − *G*_base_ versus cycle period and relaxation time. All periods divide *T* = 240 exactly, and every schedule uses a 50% duty cycle. Only period choices in the simulated grid are evaluated.

To assess sensitivity to the observation horizon, the same mean rate is evaluated for *T* = 240, 480, 960 and *T*_*c*_ = 4, 12, 30, 60, 120, with *τ*_*m*_ = 1, 10, 100. Figure S4 shows that *G* increases with the tested cycle periods across all horizons. The agreement of the rankings supports numerical robustness for these settings, not a global treatment-optimization claim.

### 5.5 Locally matched switching and fitness memory

To compare where memory enters the dynamics, we match the additional memory-induced contribution to 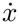 at *x*_ref_ = 1*/*2:

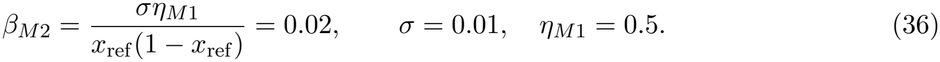

For M3 we divide that reference-state effect equally, using *η*_*M*3_ = 0.25, *β*_*M*3_ = 0.01. This matches the *memory contribution to dormant-fraction velocity*, not the complete response or fitness penalty. Figure 5 shows visually similar dormant-fraction trajectories but distinct growth outcomes because

**Figure 5:**
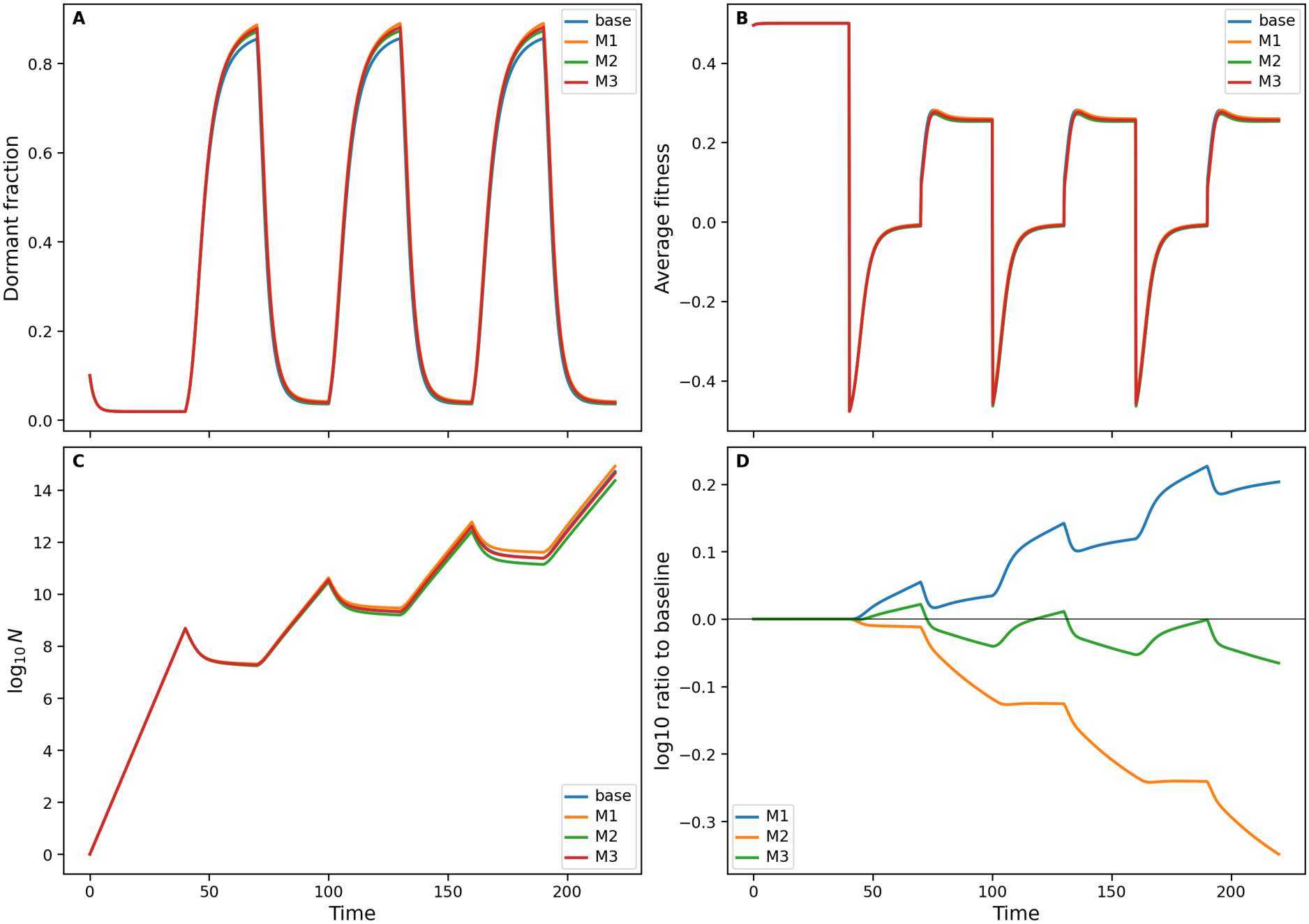
Locally matched switching- and fitness-memory feedbacks generate distinct growth responses. Under the same protocol, compare base, M1, M2 and M3 with (*η, β*) = (0.5, 0), (0, 0.02), (0.25, 0.01), *σ* = 0.01, *τ*_*m*_ = 10 and *x*_ref_ = 0.5. (A) Dormant-cell fraction; (B) mean fitness; (C) log_10_ *N*; (D) log_10_(*N*_*i*_*/N*_base_). The matching is local and does not equalize fitness.

M2/M3 modify fitness directly. The final relative changes are calculated by

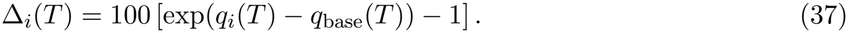

Nominal changes are +59.71% (M1), −55.21% (M2), and −13.97% (M3). The difference is not equally attributable to two independently balanced pathways: Eq. (21) and Table 2 show that the direct penalty accounts for most of the M1–M2 log-population gap. This is a consequence of the modeling hypothesis, not evidence that one biological mechanism is clinically preferable.

### 5.6 Trajectory-matched calibration and fitness-placement control

Reference-state calibration in Eq. (36) is a local construction and need not closely match the full phenotype history because *x*(*t*) may spend little time near *x*_ref_ = 0.5. To test a trajectory-level comparison we minimized

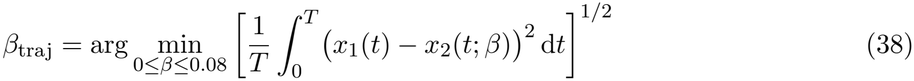

over the nominal three-cycle horizon, using uniform time sampling for the numerical objective. This gives *β*_traj_ ≈0.035997 and reduces the sampled phenotype RMS gap from 0.010977 (midpoint-matched *β* = 0.02) to 0.004206. Despite this improved phenotype agreement, log[*N*_1_(*T*)*/N*_2_(*T*)] increases from 1.271380 to approximately 1.89514. Figure 6 displays both the trajectories and the accumulated growth differences. This comparison is calibrated *to another model trajectory*, not fitted to tumor measurements.

**Figure 6:**
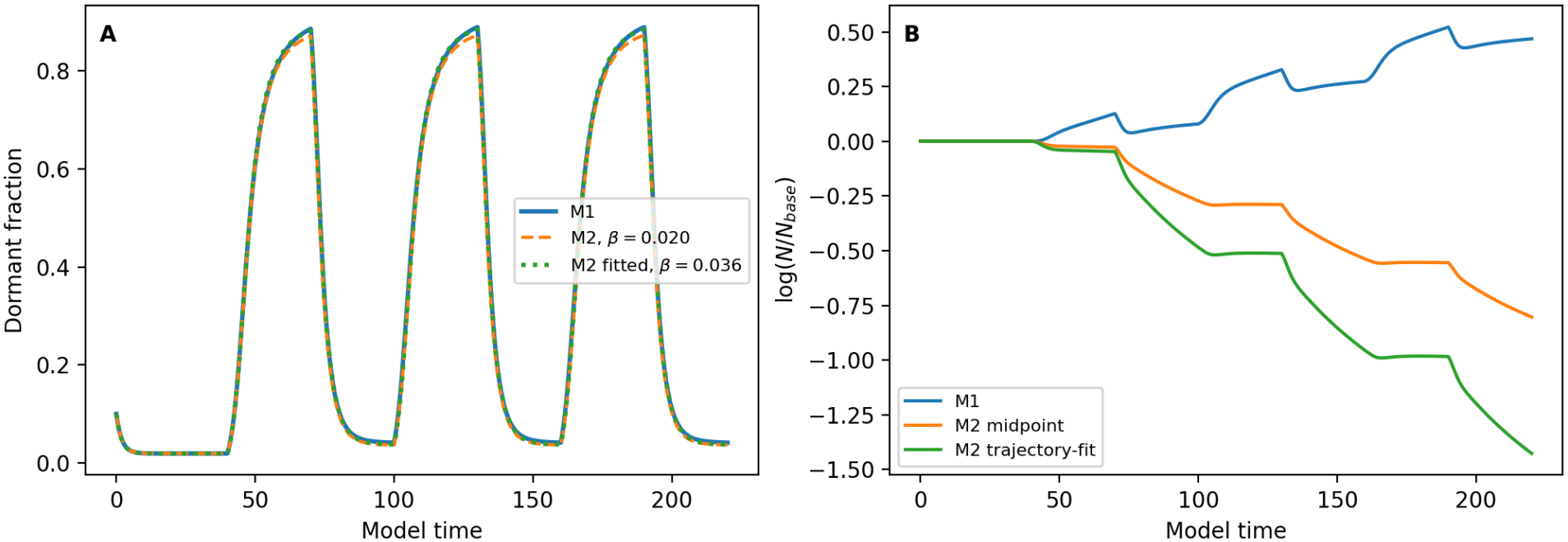
Reference-state versus trajectory-level feedback matching. (A) Dormant fractions under M1, M2 with midpoint-matched *β* = 0.02, and M2 with trajectory-fitted *β* ≈0.035997. (B) Logarithmic population ratios relative to the base model. Fitting the complete phenotype trajectory reduces its RMS mismatch while increasing separation in accumulated modeled growth.

The allocation of the same relative fitness difference is an independent structural choice. In M2, (Δ_*D*_, Δ_*P*_) = (0, −*βm*). Consider instead the dormant-fitness alternative (Δ_*D*_, Δ_*P*_) = (+*βm*, 0). Both yield the *same* fitness difference Δ_*D*_ − Δ_*P*_ = *βm*, hence identical *x*(*t*) under common initial conditions, but their mean fitnesses satisfy

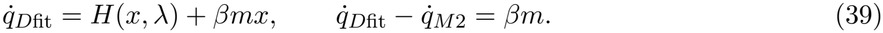

Therefore 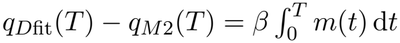 exactly. In the nominal run this integral shift is approximately 2.192886, and the dormant-fitness alternative produces a +301.37% final population change relative to the base model, versus −55.21% for M2, while the dormant fractions agree to solver tolerance (Fig. 7). This explicit counterexample shows that phenotype trajectories alone do not fix absolute fitness, and the nominal 18.9%/81.1% decomposition is specific to the chosen dormant-fitness baseline. The alternative Δ_*D*_ = +*βm* is a structural sensitivity test, not a biological claim that dormant cells grow at that rate.

**Figure 7:**
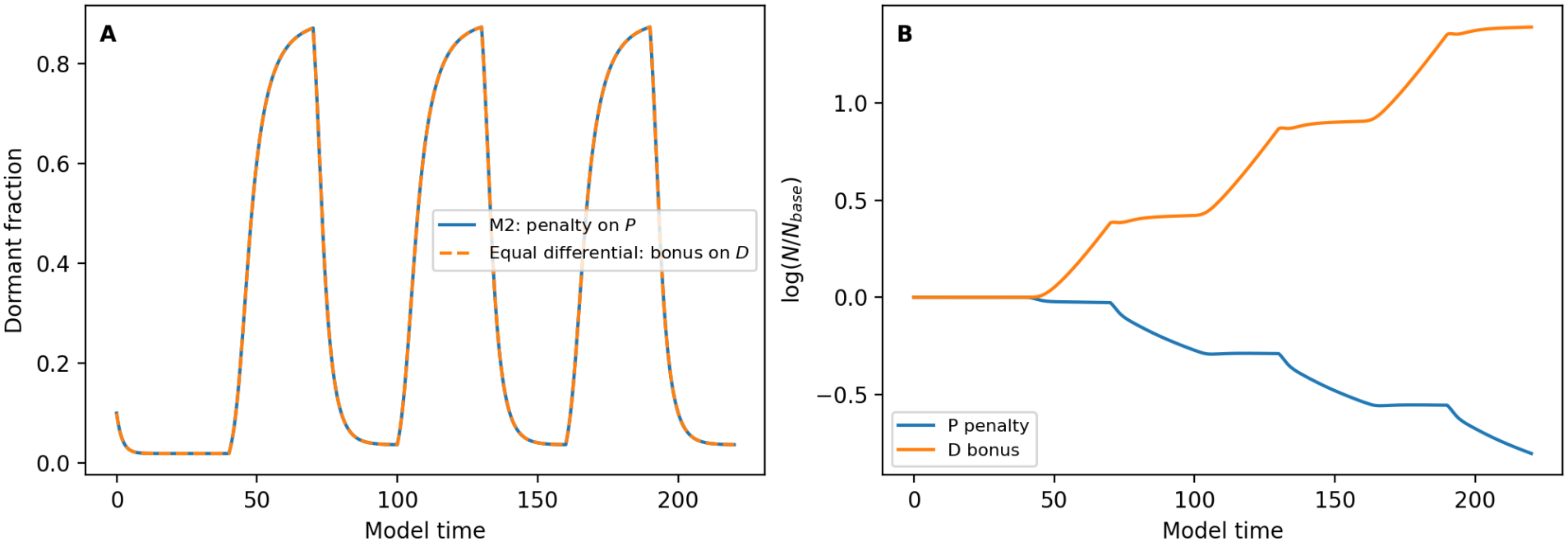
Fitness-allocation counterexample. (A) M2’s proliferating-cell penalty and an equalmagnitude dormant-cell fitness benefit produce indistinguishable phenotype trajectories because the fitness difference is identical. (B) Their cumulative modeled growth relative to the base model differs because the mean fitness depends on the absolute, not merely relative, fitness values.

### 5.7 Parameter robustness of the mechanism separation

The locally matched M2 coefficient is recalculated as *β* = *ση/*[*x*_ref_ (1 − *x*_ref_)] at each parameter point. The reference fraction is restricted to [0.25, 0.75] to avoid a small matching denominator near 0 or 1. Figure S1 varies 9 reference fractions, 9 switching strengths *η* ∈ [0.1, 0.9], and *τ*_*m*_ = 2, 10, 50 (243 comparisons). Figure S2 varies 11 reference fractions, 15 log-spaced timescales in [1, 100], and *η* = 0.2, 0.5, 0.8 (495 comparisons). In the combined 738 cases, M1 exceeds the base final modeled population and M2 lies below it. These are deterministic calculations for the fixed treatment protocol, and the non-midpoint cases are generally outside the hypothesis *β* ≤ 4*ση* of Proposition 4.1. The two full nine-panel robustness grids are consolidated into a single supplementary figure (Fig. S1) to avoid repeating the *x*_ref_-independent M1 response across heatmap rows.

To assess whether the M1-versus-base growth ordering persists beyond the original (*η, τ*_*m*_, *x*_ref_) sweeps, we performed a reproducible 2,000-point seven-dimensional Latin-hypercube exploration. The independently sampled inputs were *σ* ∈ [0.003, 0.03], *ε* ∈ [0, 0.15], *λ*_*H*_ ∈ [0.6, 1.2], *λ*_*L*_*/λ*_*H*_ ∈ [0.05, 0.6], the high-treatment duty fraction *d* ∈ [0.25, 0.75], *x*_0_ ∈ [0.2, 0.8], and *τ*_*m*_ ∈ [5, 41], with *η* = 0.5. Each experiment uses 40 untreated time units followed by three complete 60-unit cycles, with 60*d* high-treatment and 60(1 − *d*) low-treatment units per cycle; *T* = 220 is fixed, but total administered treatment cost varies. We computed Δ*G* = *G*_1_ − *G*_base_ for each parameter combination. In the seeded sample, 45*/*2,000 combinations (2.25%) had Δ*G <* 0; the remaining 1,955 yielded Δ*G >* 0. The sampled logarithmic population differences range from approximately −0.3534 to 2.3137. Of the 45 negative cases, 44 occurred with *λ*_*L*_*/λ*_*H*_ *<* 0.2 and *λ*_*H*_ *>* 0.9. An explicit in-domain example with (*σ, ε, λ*_*H*_, *λ*_*L*_*/λ*_*H*_, *d, x*_0_, *τ*_*m*_) = (0.01, 0, 1.2, 0.05, 0.5, 0.2, 10) gives log[*N*_1_(*T*)*/N*_base_(*T*)] ≈−0.3606. For the same parameters with complete withdrawal during the low phase (*λ*_*L*_ = 0, outside the stated Latin-hypercube box), we obtain log[*N*_1_(*T*)*/N*_base_(*T*)] ≈−0.4663. Hence a larger dormant fraction under M1 does *not* guarantee larger cumulative modeled growth under near-withdrawal low-treatment phases. The earlier 96-case sample detected no reversals, but that finite sample did not support a universal positive-ordering claim.

Figure 8 also reports partial rank correlation coefficients (PRCCs) computed from the same 2,000 runs, after ranking each parameter and regressing out the ranks of the other six inputs. The PRCCs are 0.770 for *σ*, −0.375 for *ε*, −0.521 for *λ*_*H*_, 0.828 for *λ*_*L*_*/λ*_*H*_, 0.401 for *d*, 0.002 for *x*_0_, and −0.067 for *τ*_*m*_. These are conditional rank associations within the specified sampling box, not variance-based Sobol indices or causal effects. Their interpretation is limited if parameter-output relations are strongly non-monotonic.

**Figure 8:**
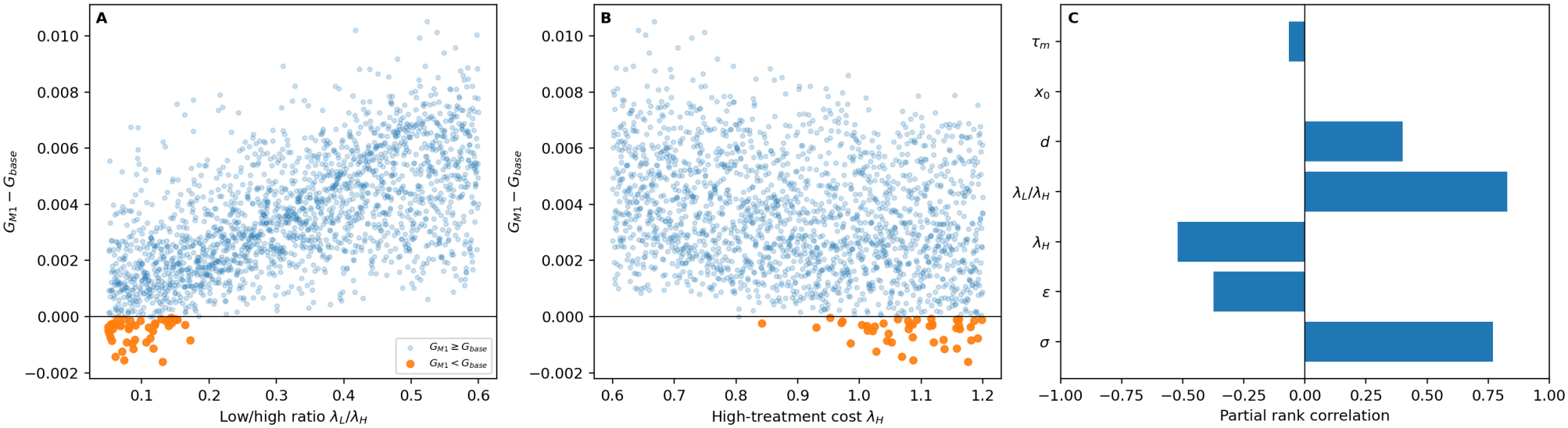
Expanded robustness of M1 relative to the base model. Two thousand seeded seven-dimensional Latin-hypercube scenarios with *T* = 220. (A) Mean logarithmic growth difference against the low-to-high treatment cost ratio and (B) against high-treatment cost; highlighted points mark the 45 observed negative differences. (C) Partial rank correlations controlling for the six remaining sampled input ranks. These results describe the tested parameter domain, not a universal treatment-ordering theorem.

The vector-field residual illustration is provided as Supplementary Fig. S3; it is a visualization of the exact algebraic identity rather than an independent identifiability experiment.

### 5.8 Adjoint verification and phase-resolved growth reversals

The exact identity in Proposition 4.3 separates the high- and low-treatment contributions to M1 relative to the base model. Under the nominal protocol the phase contributions are *C*_*H*_ = +0.622880 and *C*_*L*_ = −0.154685, producing a positive final log difference +0.468195. Under *λ*_*H*_ = 1.2, *λ*_*L*_ = 0.06, *d* = 0.5 and the same three 60-unit cycles, they are *C*_*H*_ = +0.441172 and *C*_*L*_ = −0.801796, giving a reversal of −0.360624. With nine 20-unit cycles over the same total treated horizon, the corresponding sum becomes +0.797531. The exact phase criterion shows why the outcome changes; the adjoint formula provides a complementary first-order interpretation. The first-order coefficients in Table 3 were computed by augmenting the segmented baseline ODE with cumulative sensitivity states and integrating alongside a continuous interpolant of the backward adjoint. The six representative checks illustrate the expected small-feedback accuracy, but they are not a validation of sign prediction at nominal strength. In the complete 2,000-point seven-parameter sample at *η* = 0.5, the adjoint identified only 13 of 45 exact negative M1/base cases. It missed 32 actual reversals and did not predict a reversal where the exact response was positive (Table 4). Thus first-order sensitivity is useful for local interpretation but does not classify most finite-feedback reversals in this sample. Figure 9 maps both signs across complete cycle periods dividing the 180-unit treated horizon and low/high dose ratios, holding *λ*_*H*_ = 1.2, *σ* = 0.01, *ε* = 0, *τ*_*m*_ = 10, *x*_0_ = 0.1, and *η* = 0.5 fixed. The exact and first-order boundaries differ, demonstrating the importance of nonlinear feedback at finite strength. The original adjoint time traces appear in Supplementary Fig. S5.

**Table 3:** Exact versus adjoint-predicted logarithmic growth differences. The first-order prediction is evaluated on the base trajectory. The treatment horizon is fixed at *T* = 220 and the 20-unit protocol has nine complete cycles.

| Scenario | Feedback | Exact Δ*q* | First order |
|---|---|---|---|
| Nominal M1 | *η* = 0.05 | +0.05537 | +0.05624 |
| Nominal M1 | *η* = 0.50 | +0.46820 | +0.56237 |
| Nominal M2 | *β* = 0.02 | −0.80319 | −0.81584 |
| Near withdrawal, 60-unit cycles | *η* = 0.05 | −0.01610 | −0.014532 |
| Near withdrawal, 60-unit cycles | *η* = 0.50 | −0.36062 | −0.145321 |
| Near withdrawal, 20-unit cycles | *η* = 0.50 | +0.79753 | +0.893635 |

**Table 4:** Exact versus first-order reversal classification across 2,000 parameter scenarios. Negative means log(*N*_1_(*T*)*/N*_base_(*T*)) *<* 0; the first-order column is the adjoint prediction evaluated at *η* = 0.5.

|  | Adjoint negative | Adjoint nonnegative |
|---|---|---|
| Exact negative | 13 | 32 |
| Exact nonnegative | 0 | 1955 |

**Figure 9:**
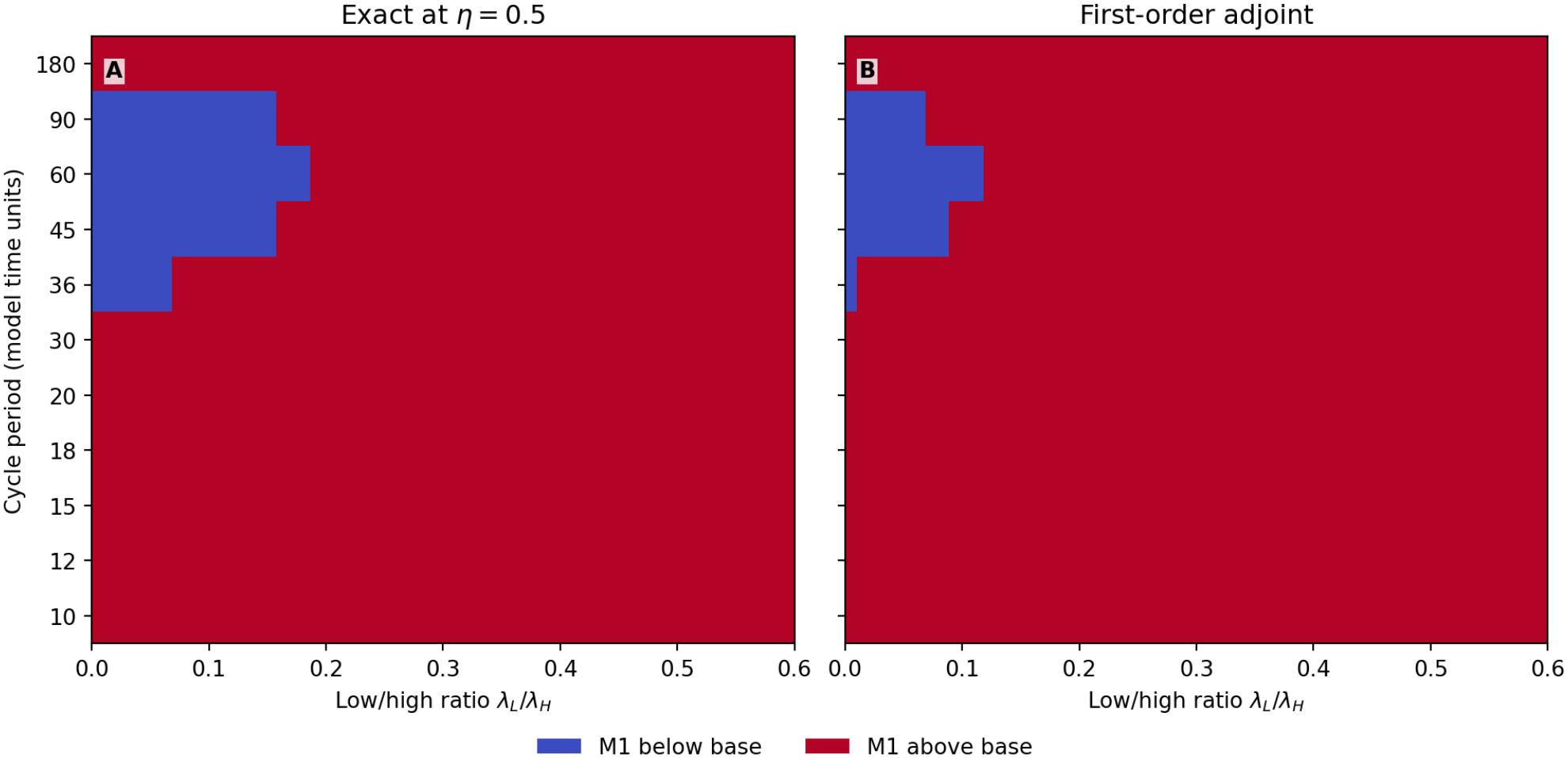
Exact versus first-order reversal map. Negative M1/base log-population differences for treatment periods dividing 180 and *λ*_*L*_*/λ*_*H*_ ∈ [0, 0.6], with a 40-unit untreated lead-in and fixed *λ*_*H*_ = 1.2. (A) Exact nonlinear response at *η* = 0.5; (B) adjoint first-order sign. The discrepancies reveal finite-feedback effects absent from linear sensitivity.

### 5.9 Density-dependent growth and time-integrated population response

The original auxiliary population equation is unbounded. As a sensitivity analysis, we added phenotype-independent density mortality to the number equation, 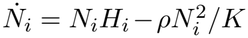, with *ρ* = 0.2 and the same unchanged phenotype equations. Because this mortality is shared by both phenotypes within a model, the dormant-fraction dynamics are unchanged. In this illustrative extension, the final log-population difference becomes nearly independent of *K* once both populations saturate. We therefore supplement the endpoint with the time-integrated difference

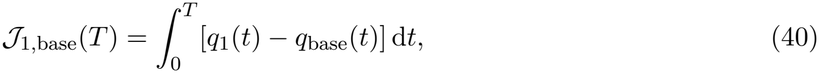

which records differences along the trajectory rather than only at *T*. This is a model-based log-population exposure measure, not a clinical endpoint. For nominal treatment, *J*_1,base_ is approximately 14.604 at *K* = 10^4^ and 14.605 at *K* = 10^6^, while the saturated endpoint log ratio is approximately 0.007102 in both cases. For the near-withdrawal protocol the integrated difference is about +3.484 at those capacities even though the unbounded population comparison gives a negative integrated difference (−25.674). Density dependence can therefore change both the magnitude and interpretation of the treatment-memory effect. Figure 10 presents these temporal measures; raw numerical outputs and the alternative fixed-budget sequence enumeration are archived as supplementary computations.

**Figure 10:**
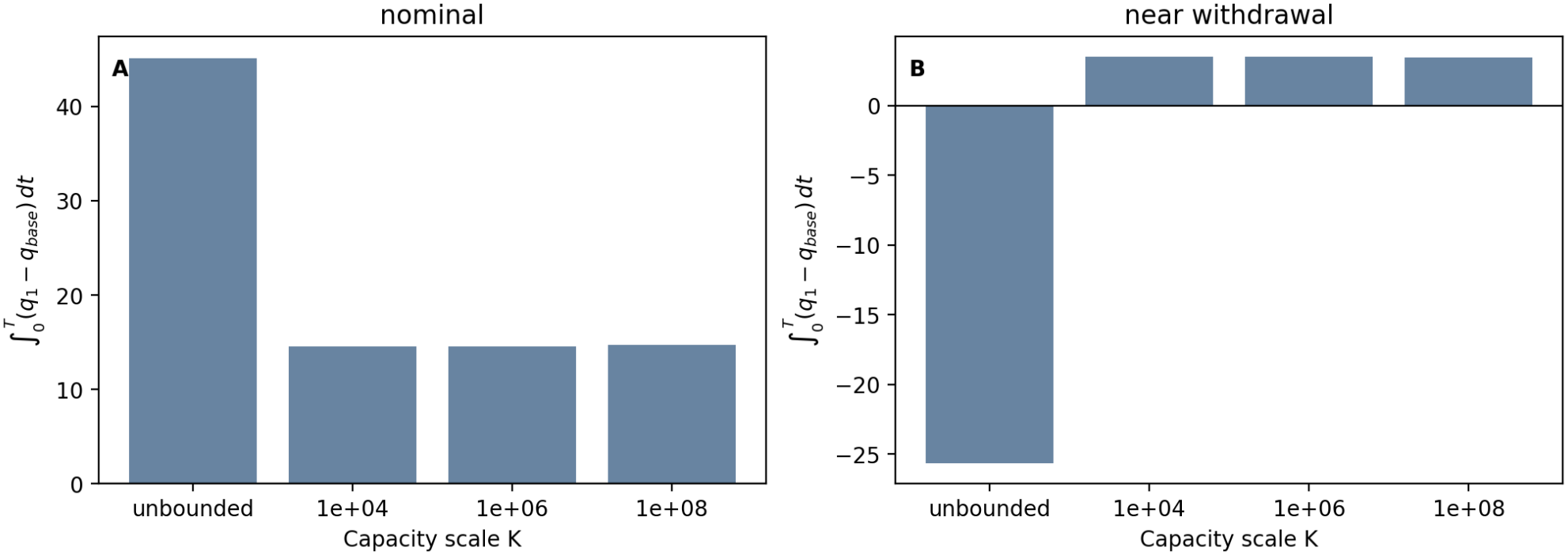
Time-integrated log-population differences under density-dependent mortality. (A) Nominal and (B) near-withdrawal treatment; each panel compares unbounded auxiliary growth with three illustrative density capacity scales. The vertical axis is J_1,base_ from Eq. (40), rather than the nearly saturated endpoint ratio.

### 5.10 Alternative switching law: dependence on feedback form

The M1 law preserves the sum of the two conversion rates. To test whether this symmetry is essential for the qualitative result, we additionally consider an *entry-only* alternative,

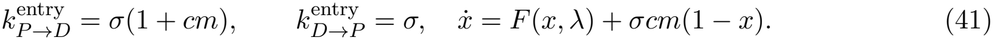

Matching its additional dormant-fraction drift to the original M1 at *x*_ref_ = 0.5 gives *c* = *η/*(1 − *x*_ref_) = 1. This alternative keeps a nonnegative entry rate and retains the same memory equation, but does not fix the sum of switching rates. Under the nominal schedule, the entry-only model yields log[*N*_entry_(*T*)*/N*_base_(*T*)] ≈0.83740 and an endpoint increase of approximately 131.03%, compared with 59.71% for M1. The two dormant-fraction trajectories differ by an unweighted sampled RMS of about 0.0133 on the plotted time grid (Figure 11). Thus the direction of the endpoint change persists in this single structural control, but its *magnitude* is sensitive to the specific switching-feedback law. One alternative is not enough to establish robustness over arbitrary admissible forms.

**Figure 11:**
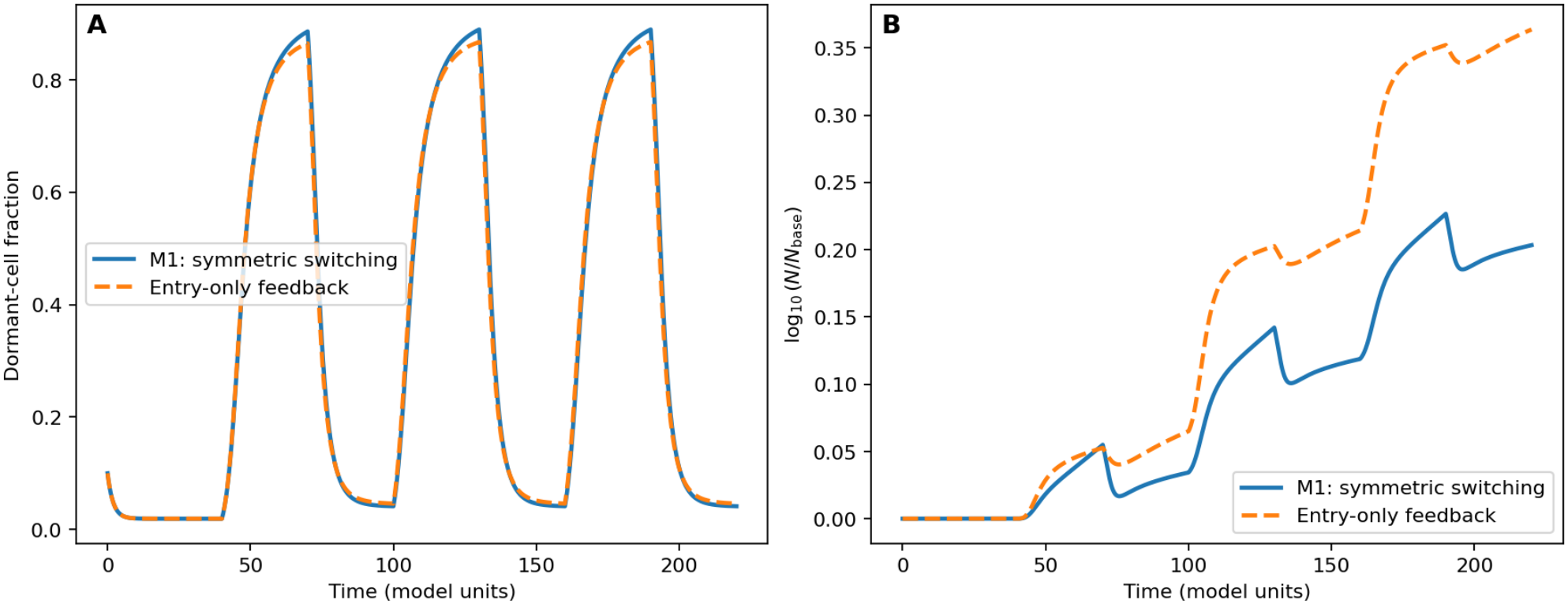
Sensitivity to the form of treatment-memory-dependent switching. Comparison of M1’s balanced bidirectional switching law (solid) with the entry-only law (dashed), locally matched at *x*_ref_ = 0.5. (A) Dormant fraction; (B) log_10_(*N/N*_base_). The entry-only law produces a larger endpoint change with otherwise common nominal settings, indicating that the magnitude of the memory effect is feedback-form dependent.

**Figure 12:**
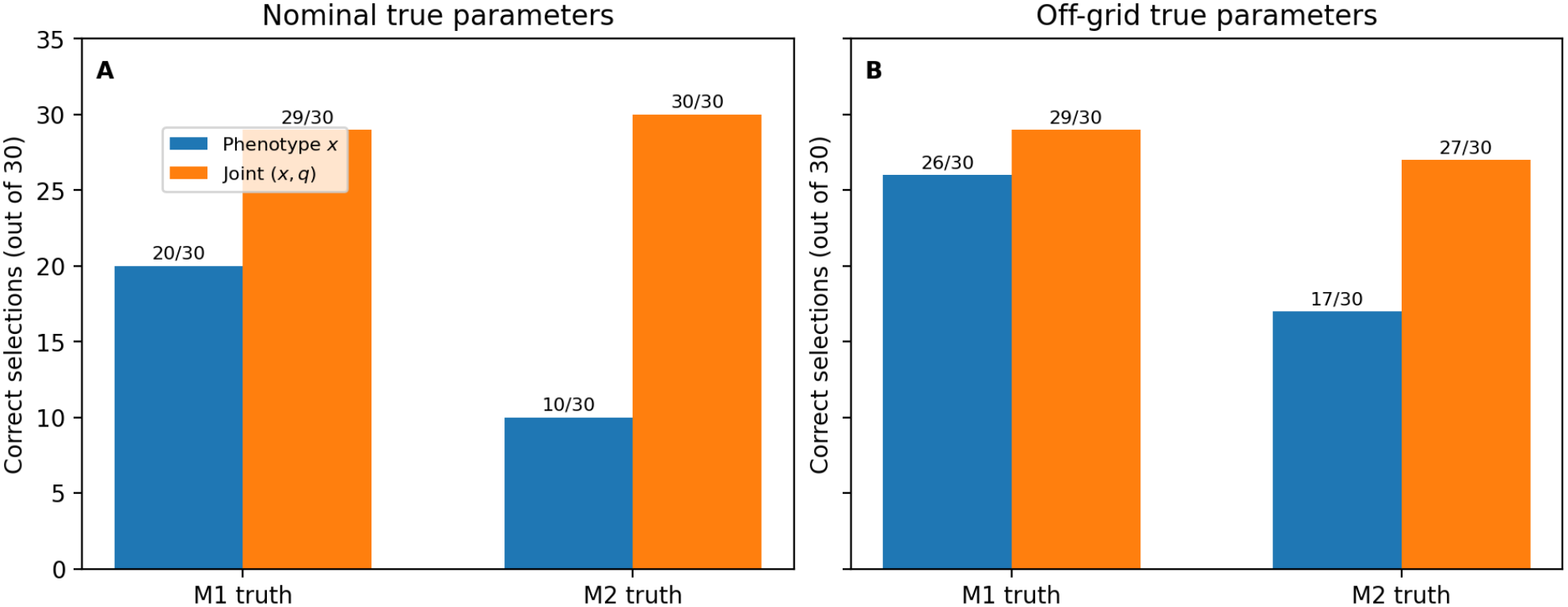
Continuous-fit discrimination with unknown baseline parameters. Correct model selections for 30 replicates per truth/model, comparing phenotype-only with joint phenotype and log-population data. Nominal and off-grid truth settings are evaluated separately; candidate feedback and baseline parameters are continuously estimated by multistart nonlinear least squares. Counts are finite-sample, in-sample comparisons.

### 5.11 Exploratory Model Selection from Noisy Synthetic Observations

As a separate test of baseline uncertainty, we fitted *σ, ε*, and the relevant feedback coefficient simultaneously by multistart continuous least squares, rather than by a discrete parameter grid. We considered both nominal truths (*σ, ε, η, β*) = (0.01, 0, 0.5, 0.02) and off-grid truths with *σ* = 0.0115, *ε* = 0.03, *η* = 0.53 (M1), or *β* = 0.023 (M2). For each mechanism and truth setting, 30 independent synthetic datasets were generated with the same 22 measurement times and Gaussian noise levels as in the fixed-baseline experiment. Each model was optimized from two initial guesses within *σ* ∈ [0.002, 0.05], *ε* ∈ [0, 0.2], *η* ∈ [0, 1] (M1) or *β* ∈ [0, 0.08] (M2). With joint (*x, q*) observations, correct counts were 29/30 and 30/30 for nominal M1 and M2, and 29/30 and 27/30 for off-grid M1 and M2. Phenotype-only counts were respectively 20/30, 10/30, 26/30, and 17/30. We report signed 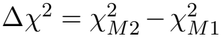 (positive favors M1), not absolute gaps. These limited in-sample counts demonstrate that grid alignment can inflate discrimination claims; they are not estimates of general clinical identifiability. Continuous fits and per-dataset signed losses are supplied in the reproducibility data.

## 6 Discussion and Conclusions

We developed and analyzed three treatment-history-dependent extensions of a two-phenotype evolutionary model of glioma dormancy, distinguishing memory effects on phenotype switching (M1), proliferative fitness (M2), and their combination (M3). The central objective was to determine how the placement of treatment memory influences dormant-cell fractions and cumulative population growth under identical treatment histories. Our results demonstrate that memory-dependent mechanisms with locally matched contributions to phenotype dynamics can nevertheless produce substantially different population-growth predictions. This distinction highlights the importance of identifying how treatment history influences cellular dynamics, rather than considering memory solely as an additional state variable.

The analytical results provide a mathematical framework for interpreting these differences. The biologically admissible state region is invariant, and the equilibrium and local stability analysis characterizes the response to constant treatment. The phenotype comparison establishes an ordering between M1 and the base model and, under additional parameter restrictions, between M1 and M2.

More importantly, the exact logarithmic population-growth identity separates the M1–M2 difference into contributions from phenotype composition and the direct fitness-memory penalty. Under the nominal three-cycle protocol, approximately 18.9% of the logarithmic population difference is associated with phenotype composition, while 81.1% is associated with the explicit fitness-memory term in M2. Thus, the substantial difference in predicted population growth is driven primarily by the structural placement of memory in the fitness equation, rather than by differences in the dormant-cell fractions alone. These percentages describe an exact decomposition within the specified mathematical models and should not be interpreted as experimentally established causal contributions.

A further analytical result identifies distinct dynamical signatures of switching-memory and fitness-memory feedback. By combining the residuals of the phenotype-fraction and logarithmic population-growth equations, the contributions of the two memory mechanisms can be separated algebraically when the base-model parameters and memory trajectory are known and memory does not alter dormant fitness. This provides a potential connection between the mathematical formulation and experimental mechanism identification. Preliminary synthetic-data experiments support this interpretation: under the investigated observation and noise settings, jointly using phenotype fractions and population growth improved model-selection accuracy relative to using phenotype fractions alone. Nevertheless, this structural separation does not establish practical identifiability under realistic experimental conditions, where observations may be sparse or noisy and the underlying parameters may be uncertain.

The numerical experiments further demonstrate that the predicted population response depends on the strength, duration, and placement of treatment memory. Under the nominal three-cycle protocol, the targeted sweeps over *η, τ*_*m*_, and *x*_ref_ gave M1 growth above and M2 growth below the base model. In contrast, the broader seven-parameter sample includes negative M1 growth differences under strong-dose, near-withdrawal schedules. Consequently, the earlier sign ordering is specific to the nominal protocol and cannot be extrapolated to other treatment schedules. An alternative entry-only switching formulation preserved the direction of the M1 population response while substantially altering its magnitude, demonstrating that quantitative predictions remain sensitive to the assumed feedback law.

Two additional tests sharpen the interpretation. Full-trajectory matching reduces the M1–M2 phenotype discrepancy but increases their cumulative-growth difference, showing that the latter is not explained solely by midpoint-calibration error. Conversely, allocating an equivalent memory-dependent fitness difference to dormant rather than proliferating fitness leaves the phenotype trajectory invariant while reversing the sign of the growth comparison. Hence the contrast is not robust to arbitrary redistributions of absolute net growth between phenotypes, and the algebraic residual separation is only identifiable under the declared Δ_*D*_ = 0 assumption. In an expanded seven-parameter sample, the M1-versus-base growth difference changed sign in low-dose-holiday-like regimes with high treatment costs. The nominal increase is therefore protocol-dependent rather than a robust inequality over the sampling domain.

The stronger schedule-dependent conclusion is that switching memory can either raise or lower modeled growth under the assumed feedback laws. The exact phase-integral result explains the reversal, whereas the adjoint characterizes only the infinitesimal-feedback response and misses many reversals at nominal switching strength. Both conclusions remain sensitive to how net fitness is assigned to the two phenotypes. The density-dependent control also shows that saturation can substantially reduce the endpoint differences. The treatment-frequency experiments provide additional information about the interaction between memory timescales and treatment scheduling. For the investigated equal-exposure high/low protocols, shorter treatment cycles produced lower finite-horizon mean logarithmic growth in both the base and memory-dependent models. At the nominal high/low treatment levels tested in the frequency and horizon comparisons, the memory extensions preserved the observed qualitative period ordering. This numerical ordering does not extend to other dose choices: the finite-feedback reversal map in Fig. 9 shows that changing the cycle period can change the sign of the M1/base growth difference. The analytical periodic-response and averaging results clarify the role of memory filtering and rapidly alternating exposure, but they do not establish globally optimal treatment schedules or long-time population ordering.

Several modeling assumptions limit the biological interpretation of these findings. The framework represents the tumor population using two aggregate phenotypes and a single treatment-memory variable shared across cells. It does not explicitly account for spatial heterogeneity, resource limitations, pharmacokinetics, or interactions with the tumor microenvironment. The symmetric modification of phenotype-switching rates in M1 and the residual fitness penalty in M2 are phenomenological representations of possible treatment-history effects. In particular, persistent fitness suppression represents only one direction of treatment-associated adaptation; acquired tolerance or resistance may require alternative feedback signs or functional forms. Consequently, the conclusions depend on the stated assumptions and should not be interpreted as universal biological responses.

An additional limitation concerns the interpretation of total population growth. The auxiliary growth equation assumes that mean evolutionary fitness represents an effective per-capita net growth rate. Because the model does not include density-dependent saturation or a calibrated physical timescale, the absolute magnitude of the modeled population cannot be interpreted directly as tumor cell count or physical tumor volume. Relative population comparisons and finite-horizon mean logarithmic growth rates therefore provide more appropriate measures of the modeled response. For perspective, the nominal +59.71% endpoint M1/base ratio corresponds to *G*_1_ − *G*_base_ ≈0.002128, only about 1.38% of the nominal base rate. The density-dependent sensitivity calculation in Section 5.9 shows that endpoint ratios can be strongly attenuated by saturation. Likewise, the nominal switching, memory, and fitness parameters are illustrative rather than experimentally estimated, and the predicted reactivation delays and population differences should not be interpreted as quantitative clinical predictions.

Future investigations should prioritize connecting the proposed feedback mechanisms to longitudinal glioblastoma measurements. In particular, simultaneous observations of phenotype fractions and total viable-cell populations, including measurements during treatment withdrawal, could help assess whether switching-memory and fitness-memory effects are distinguishable in practice. Further analysis of sparse and noisy observations, uncertain baseline parameters, and alternative admissible feedback functions would clarify the robustness and identifiability of the proposed mechanisms. On the mathematical side, sharper population-comparison conditions or validated numerical bounds could strengthen the connection between the analytical results and the nominal numerical experiments.

Overall, this study shows that the consequences of treatment history depend not only on the persistence of memory but also on the biological mechanism through which it acts. The exact population-growth decomposition and the analytical separation of memory-feedback signatures provide complementary ways to distinguish phenotype-switching effects from direct fitness effects. Together with the numerical comparisons, these results establish a framework for formulating and evaluating mechanism-specific hypotheses about treatment-associated glioma dormancy, while identifying the experimental measurements needed to assess their biological relevance.

## Acknowledgments

G.F.-J. acknowledges support from the Undergraduate Research Opportunity Program (UROP) at the University of Michigan. N.A. acknowledges support from NIH/NCCIH grant R01AT013188 and NSF DMS grant #2619580.

## Code and Reproducibility

All numerical simulations were performed in Python using SciPy’s ODE solvers, primarily LSODA, with additional numerical verification using DOP853. The complete model implementation, figure-generation scripts, and numerical analysis codes will be made publicly available upon acceptance of the manuscript through our GitHub repository.

## A Supplementary Material

### A.1 Consolidated parameter maps and ideal residual illustration

The two robustness-grid experiments are presented as consecutive supplementary figures to preserve complete numerical context without presenting repeated *x*_ref_-independent M1 rows in the main figures. The upper grid varies switching strength and reference fraction at fixed memory timescales; the lower grid varies timescale and reference fraction at fixed switching strengths. The repeated vertical structure of M1 is expected because its governing equation contains no matching reference fraction.

### A.2 Adjoint, density-dependent, and data-anchoring limitations

The accompanying archive contains the numerical adjoint checks, phase-resolved growth contributions, density-dependent trajectory integrals, and the fixed-budget schedule enumeration as separate CSV files. A random 10,000-case numerical scan of stable equilibria over the averaged-dose ranges generated by the seven-parameter sampling box found a positive value of for all tested draws. This is not a proof over the continuous parameter set and does not 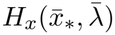 imply positivity at finite cycle periods or finite *η*. We retain the exact phase criterion as the nonperturbative statement. Larsson et al. [21] provide public longitudinal glioblastoma cell-state data and code, but their cell-state model has multiple transcriptional states and distinct measured growth rates. Mapping those observations to the two dormant/proliferating compartments and estimating the treatment-memory timescale requires new biological assumptions; we therefore do not present the illustrative *σ* = 0.01 or the drug-holiday pattern as experimentally calibrated. Public datasets and software can be accessed through the Larsson et al. study and its linked STAG repository; this study does not claim a fit of those datasets to the present two-phenotype model.

**Additional numerical illustrations**.

### A.3 Numerical examination of constant-treatment equilibria

To complement the equilibrium existence and local stability analysis in Section 3, we numerically examined the number of biologically admissible equilibria under constant treatment.

For each model, the equilibrium memory state is *m*_*\**_ = *λ*_0_*/λ*_*H*_, and the equilibrium dormant fraction satisfies

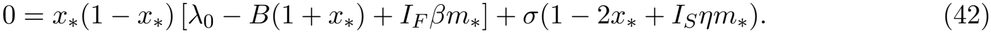

Here, (*I*_*S*_, *I*_*F*_) = (0, 0), (1, 0), (0, 1), and (1, 1) correspond to the base model, M1, M2, and M3, respectively.

Using the nominal parameter values specified in Table 5, with all parameters except *λ*_0_ held fixed, we computed the roots of Eq. (42) at 1,001 equally spaced treatment levels *λ*_0_ ∈ [0, 1]. Only real roots in the biologically admissible interval [0, 1] were counted.

**Table 5:**
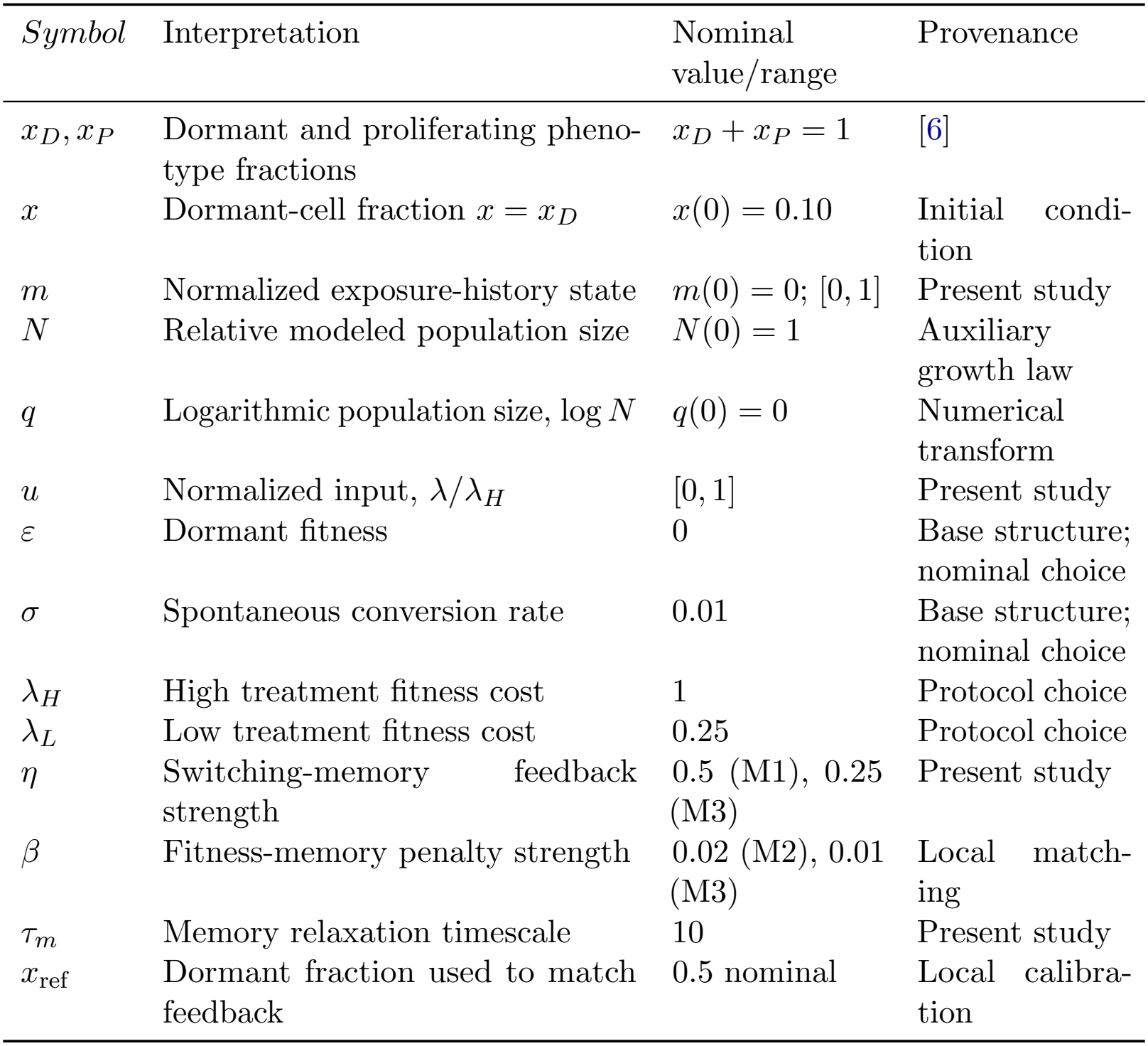
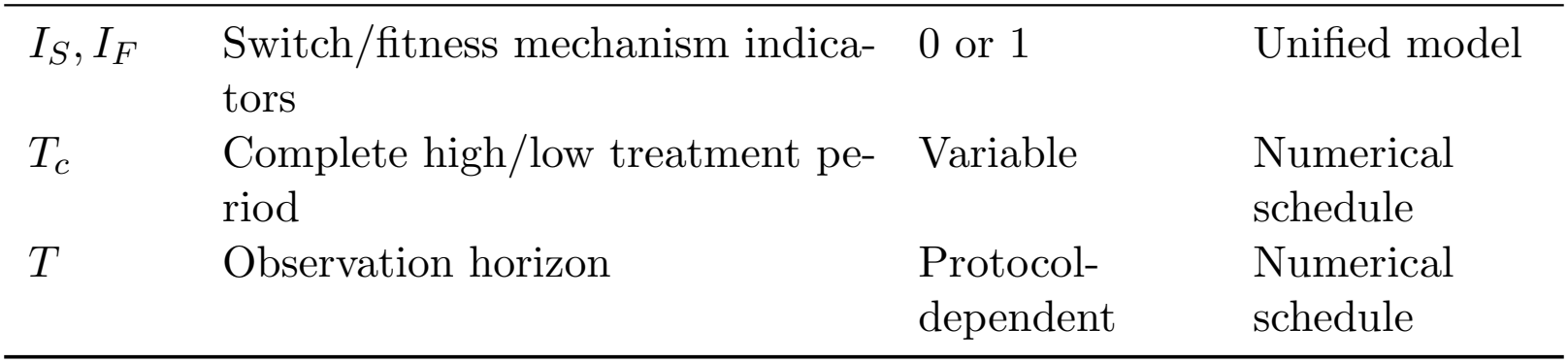
Model states, parameters, and nominal comparison values.

| *Symbol* | Interpretation | Nominal value/range | Provenance |
|---|---|---|---|
| x<sub>D</sub>, x<sub>P</sub> | Dormant and proliferating pheno-type fractions | x<sub>D</sub> + x<sub>P</sub> = 1 | [6] |
| x | Dormant-cell fraction x = x<sub>D</sub> | x(0) = 0.10 | Initial condi-tion |
| m | Normalized exposure-history state | m(0) = 0; [0, 1] | Present study |
| N | Relative modeled population size | N(0) = 1 | Auxiliary growth law |
| q | Logarithmic population size, log N | q(0) = 0 | Numerical transform |
| u | Normalized input, λ/λ<sub>H</sub> | [0, 1] | Present study |
| ε | Dormant fitness | 0 | Base structure; nominal choice |
| σ | Spontaneous conversion rate | 0.01 | Base structure; nominal choice |
| λ<sub>H</sub> | High treatment fitness cost | 1 | Protocol choice |
| λ<sub>L</sub> | Low treatment fitness cost | 0.25 | Protocol choice |
| η | Switching-memory feedback strength | 0.5 (M1), 0.25 (M3) | Present study |
| β | Fitness-memory penalty strength | 0.02 (M2), 0.01 (M3) | Local match-ing |
| τ<sub>m</sub> | Memory relaxation timescale | 10 | Present study |
| x<sub>ref</sub> | Dormant fraction used to match feedback | 0.5 nominal | Local calibra-tion |
| I<sub>S</sub>, I<sub>F</sub> | Switch/fitness mechanism indica-tors | 0 or 1 | Unified model |
| T<sub>c</sub> | Complete high/low treatment pe-riod | Variable | Numerical schedule |
| T | Observation horizon | Protocol-dependent | Numerical schedule |

**Table 6:**
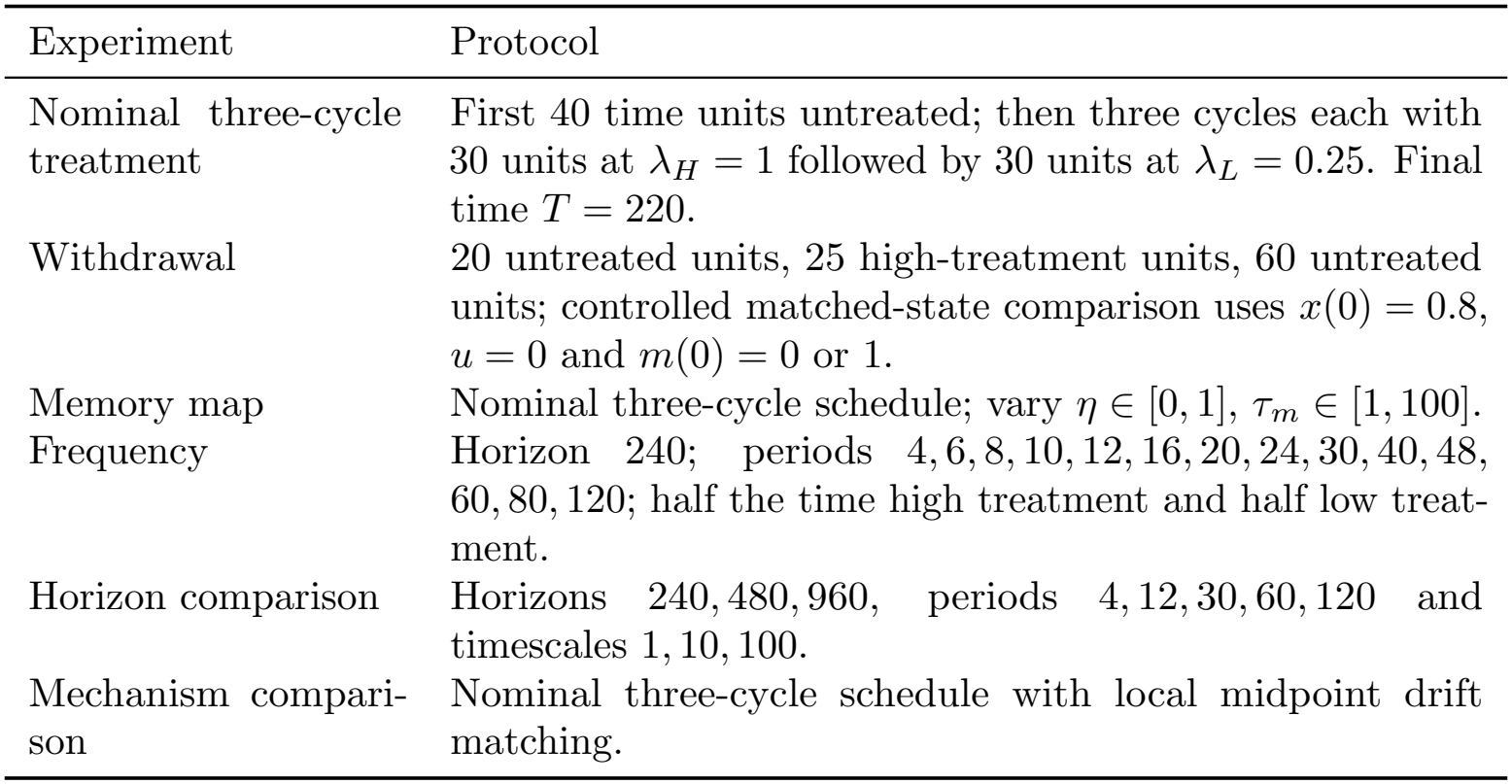
Treatment protocols for the main numerical comparisons.

| Experiment | Protocol |
|---|---|
| Nominal three-cycle treatment | First 40 time units untreated; then three cycles each with 30 units at λ<sub>H</sub> = 1 followed by 30 units at λ<sub>L</sub> = 0.25. Final time *T* = 220. |
| Withdrawal | 20 untreated units, 25 high-treatment units, 60 untreated units; controlled matched-state comparison uses *x*(0) = 0.8, *u* = 0 and *m*(0) = 0 or 1. |
| Memory map | Nominal three-cycle schedule; vary η ∈ [0, 1], τ<sub>*m*</sub> ∈ [1, 100]. |
| Frequency | Horizon 240; periods 4, 6, 8, 10, 12, 16, 20, 24, 30, 40, 48, 60, 80, 120; half the time high treatment and half low treat-ment. |
| Horizon comparison | Horizons 240, 480, 960, periods 4, 12, 30, 60, 120 and timescales 1, 10, 100. |
| Mechanism compari-son | Nominal three-cycle schedule with local midpoint drift matching. |

**Table 7:** Solver-consistency summary for 36 numerical comparisons.

| Output | Methods compared | Maximum endpoint difference |
|---|---|---|
| *x*(*T*) | LSODA / DOP853 | 5.7 × 10<sup>−11</sup> |
| *q*(*T*) | LSODA / DOP853 | 1.18 × 10<sup>−7</sup> |

**Table 8:**
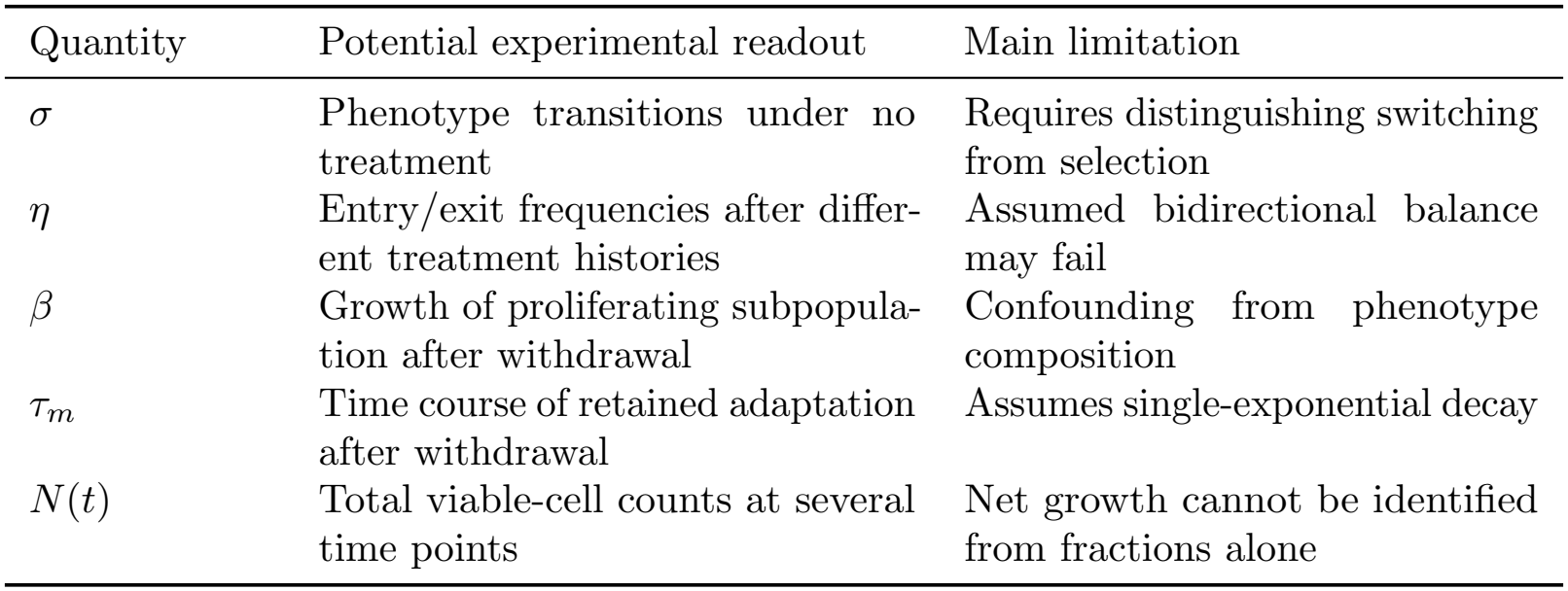
Measurements needed for subsequent parameter estimation and feedback discrimination.

| Quantity | Potential experimental readout | Main limitation |
|---|---|---|
| σ | Phenotype transitions under no treatment | Requires distinguishing switching from selection |
| η | Entry/exit frequencies after differ-ent treatment histories | Assumed bidirectional balance may fail |
| β | Growth of proliferating subpopula-tion after withdrawal | Confounding from phenotype composition |
| τ<sub>m</sub> | Time course of retained adaptation after withdrawal | Assumes single-exponential decay |
| N(t) | Total viable-cell counts at several time points | Net growth cannot be identified from fractions alone |

For all four models, exactly one admissible equilibrium was identified at each of the 1,001 sampled treatment levels. Thus, no evidence of multiple admissible equilibria was found for the nominal parameter values over the sampled treatment grid. This finite numerical examination does not constitute a proof of equilibrium uniqueness over the continuous treatment interval or for other parameter combinations.

### A.4 Model variables, nominal values, and provenance

The following values are illustrative model inputs rather than patient-level fits. The base-model structure is inherited from Ref. [6]; values and memory feedback strengths used in the present comparisons are protocol-dependent modeling choices.

**Figure S1:**
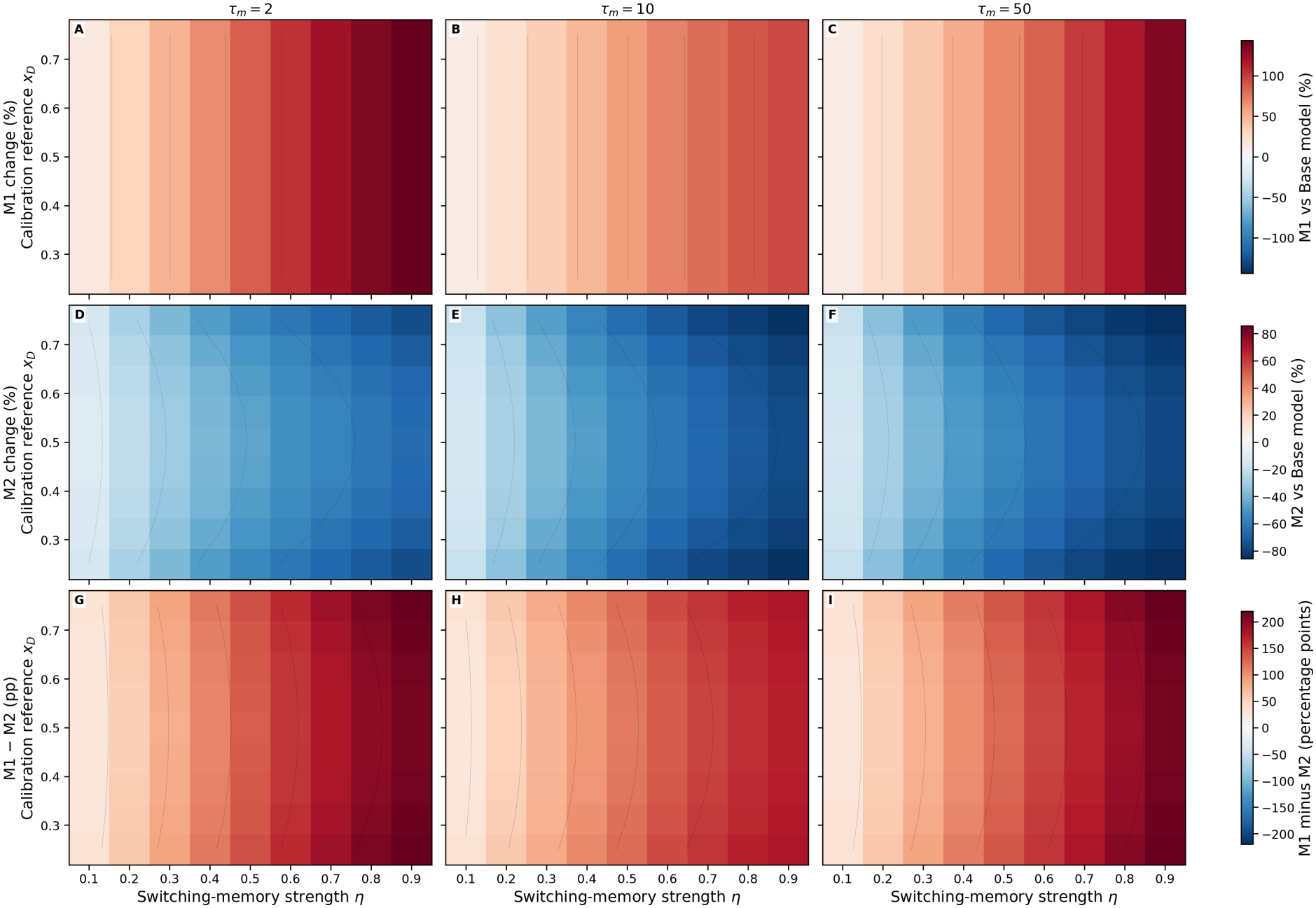
Switching strength and reference-fraction robustness. Results for 243 combinations of *η, x*_ref_, and *τ*_*m*_ = 2, 10, 50: M1 and M2 final changes relative to the base model, and their percentage-point difference.

**Figure S2:**
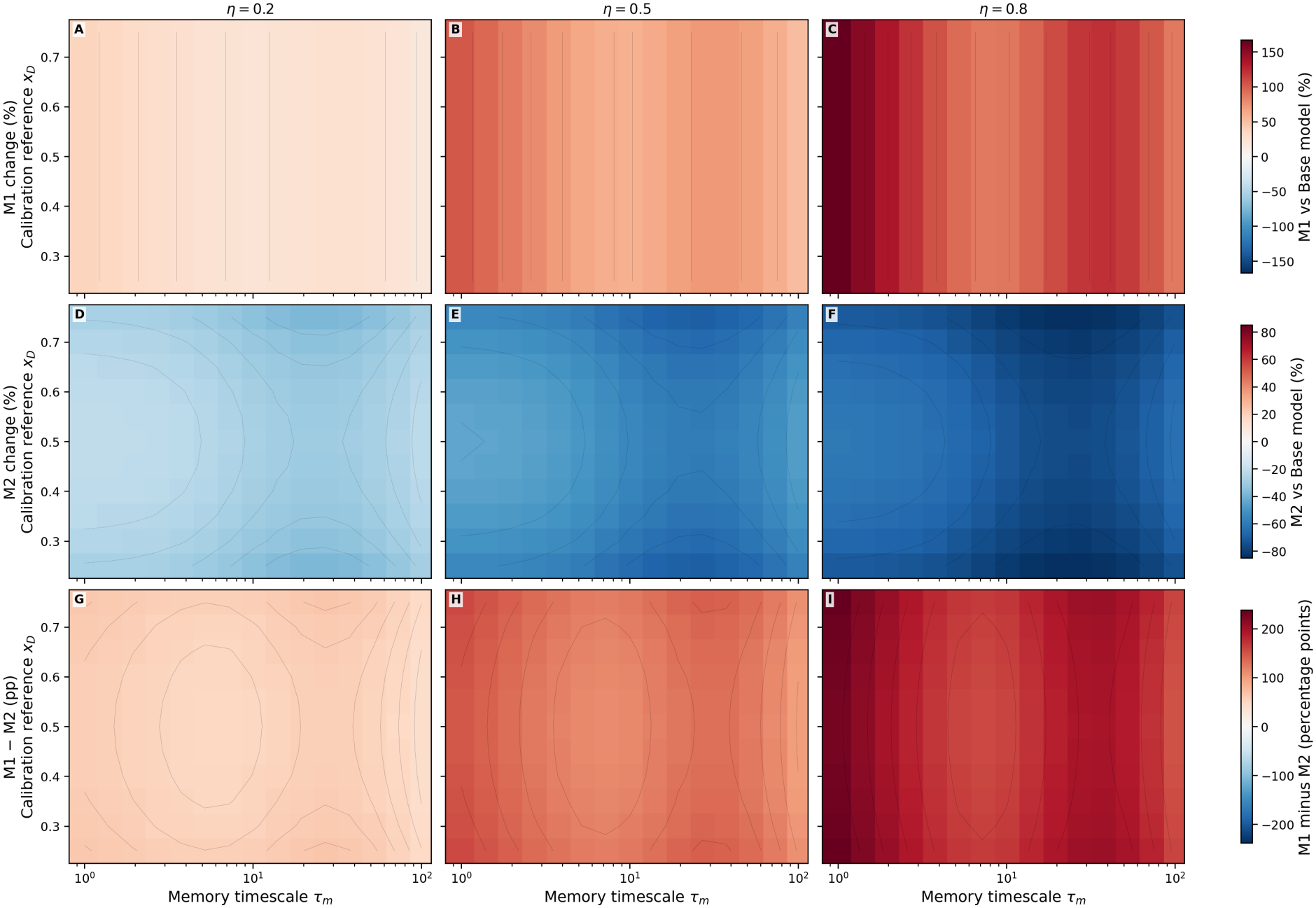
Memory timescale and reference-fraction robustness. Results for 495 combinations of *τ*_*m*_, *x*_ref_, and *η* = 0.2, 0.5, 0.8: M1 and M2 final changes relative to the base model, and their percentage-point difference.

**Figure S3:**
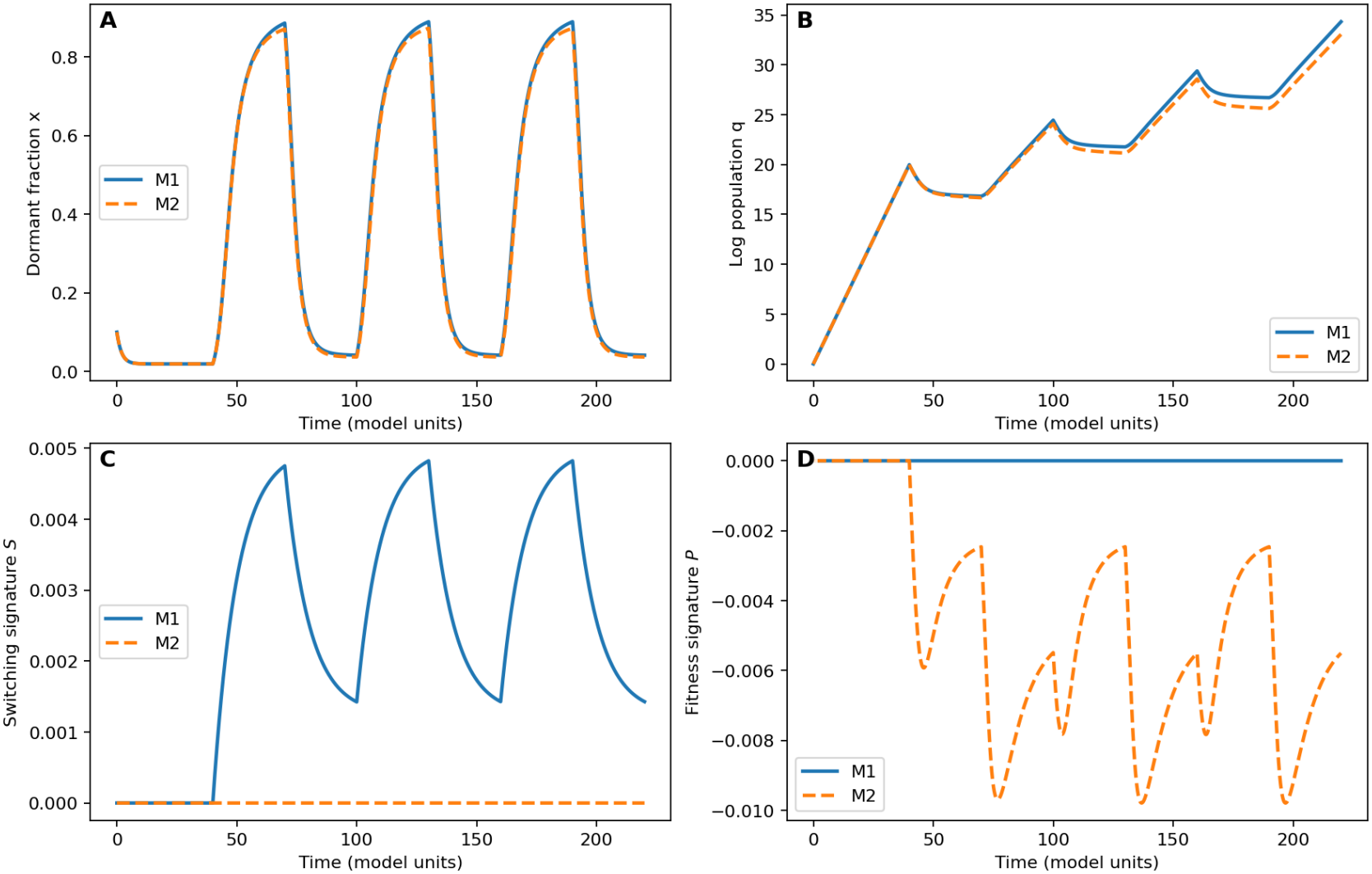
Ideal vector-field signatures. (A) Dormant fractions, (B) log modeled populations, (C) switching residual, and (D) fitness residual for M1 and M2. Residuals use derivatives from the exact model vector fields and do not demonstrate extraction from noisy experiments.

**Figure S4:**
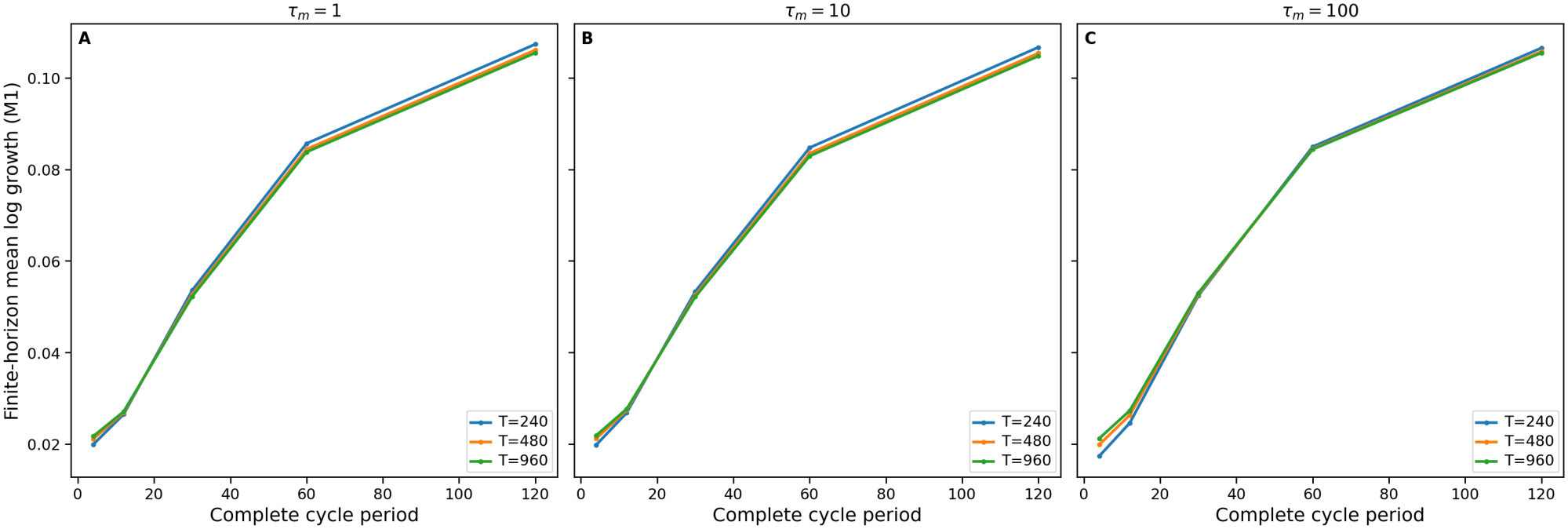
Treatment-period ordering across three observation horizons. M1 mean logarithmic growth for horizons *T* = 240, 480, 960 and periods *T*_*c*_ ∈ {4, 12, 30, 60, 120} with equal high/low exposure time. Panels (A)–(C) correspond to *τ*_*m*_ = 1, 10, 100. The displayed ordering persists within the tested grid.

**Figure S5:**
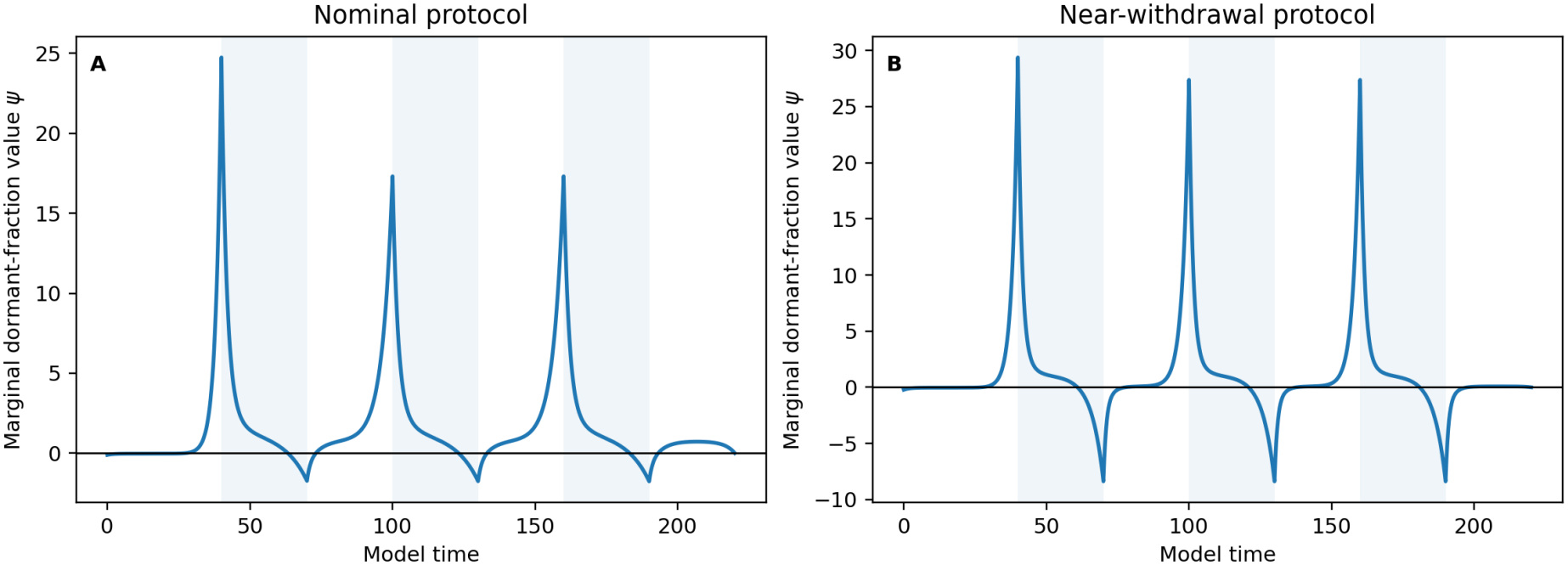
Adjoint value of dormancy along the base dynamics. (A) Nominal alternating high/low protocol; (B) strong-treatment, near-withdrawal protocol. Shading indicates high-treatment phases. Negative *ψ* identifies times when an additional small dormant-fraction perturbation reduces final modeled logarithmic growth, conditional on the subsequent schedule.

### A.5 Treatment schedules

### A.6 Numerical integration and reliability checks

The numerical package uses `scipy.integrate.solve_ivp` with LSODA and integration segmented at all treatment switches. Its default parameter core uses `rtol=3e-9, atol=1e-11`; the supplementary analyses use DOP853 for additional feedback and adjoint checks with stricter tolerances. These procedures ensure that the discontinuous treatment input is applied at the intended times. The solver-consistency audit independently compared LSODA and DOP853 on 36 selected cases, reporting maximum absolute differences of approximately 5.7×10^−11^ in endpoint dormant fraction and 1.18×10^−7^ in endpoint log population. This is a solver-consistency check, not a validated enclosure of all model solutions. data/alternative `switching_comparison.csv` documents the alternative-feedback endpoint comparison.

### A.7 Reproducibility of supplementary numerical experiments

The supplied Python scripts reproduce the figure calculations, seven-parameter sensitivity experiment, trajectory fitting, and synthetic model-selection tests. Reported trajectory-fit RMS values use 441 equally spaced points on [0, 220]; the expanded 2,000-point Latin-hypercube exploration uses seed 20261116 with independent uniform marginals over the seven explicitly stated parameter intervals; its PRCC values are calculated from the saved sampled data. The v2 unknown-baseline analysis uses multistart continuous fitting on 30 datasets per model and truth setting, including off-grid truths; the supplied numerical analysis uses continuously fitted baseline parameters. Earlier fixed-baseline, grid-fitted synthetic results are retained only as historical computational output; the manuscript’s updated unknown-baseline comparison uses continuous multistart fitting. The tabulated CSV files are supplied in the data/ folder, including signed fitting losses and exact/adjoint sign comparisons. These are numerical sensitivity and model-selection checks, not calibrated GBM parameter estimates.

### A.8 Observable quantities and future estimation experiments

For direct phenotype and cell-count measurements, observable candidates include *x*_*D*_(*t*), *x*_*P*_ (*t*) = 1 − *x*_*D*_(*t*), and relative viable-cell count *N* (*t*)*/N* (0). The residual-based structural signature result assumes known baseline parameters and direct access to 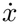 and 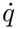; it is not automatically usable on noisy sampled trajectories.

